# The buried S2 apex of SARS-CoV-2 spike elicits an immunodominant germline-restricted public antibody response

**DOI:** 10.64898/2026.02.18.706653

**Authors:** Suncheol Park, Jacob Mischka, Nisreen Okba, Anass Abbad, Meng Yuan, Komal Srivastava, Charles Gleason, Lubbertus C.F. Mulder, Jeffrey Copps, Katrina Saam, Sandhya Bangaru, Florian Krammer, Ian A. Wilson, Viviana Simon, Andrew B. Ward

## Abstract

The continued mutational pressure on the SARS-CoV-2 S1 subunit underscores the need to target the conserved S2 region for pan-coronavirus vaccine development. A detailed molecular understanding of S2-directed immune responses is therefore essential. In this study, we identified the S2 apex as the most immunodominant epitope within the S2 subunit, eliciting robust antibody responses despite occlusion by S1, using electron-microscopy-based polyclonal epitope mapping (EMPEM) of plasma from infected and vaccinated individuals. Structure-guided sequence analysis with antibody databases revealed that antibodies targeting a poorly characterized S2 Apex-B site form a convergent public clonotype, which is predominantly derived from the IGHV3-30 germline with a 14-residue CDRH3 containing a G/S-G-S/N-Y motif. This clonotype is extensively expanded, accounting for up to 40% of total spike-reactive antibody sequence counts in individual vaccinated donors. This study elucidates the molecular basis the high-frequency elicitation of this non-neutralizing clonotype emphasizing that its immunodominance acts as a primary hurdle for universal coronavirus vaccines and underscore the need for precision antigen design to redirect immunity toward more potent neutralizing targets.

## INTRODUCTION

The coronavirus disease 2019 (COVID-19) pandemic has accelerated the development of vaccines and antibody therapeutics targeting the spike protein at an unprecedented pace^1^. The spike protein is comprised of S1 and S2 subunits based on the furin cleavage site, with S1 located upstream and S2 located downstream of this cleavage site **(Figure 1A)**. Vaccine and therapeutic antibody development has predominantly targeted the S1 subunit, where the receptor-binding domain (RBD) and N-terminal domain (NTD) are located. Although antibodies targeting the S1 region initially exhibit neutralizing activity, their effectiveness has rapidly declined as ongoing viral evolution has imposed intense mutational pressure on this region, limiting the durability of such approaches^2–6^. Thus, COVID-19 vaccine formulations have been iteratively updated to keep up with the ever changing severe acute respiratory syndrome coronavirus 2 (SARS-CoV-2) variants. Consequently, recent studies have begun to focus on the S2 subunit, a region highly conserved across coronavirus subfamilies and SARS-CoV-2 variants, emerging as a crucial alternative for pan-coronavirus immunization strategies. Recent vaccine approaches utilizing the S2 subunit have demonstrated cross-protective effects against multiple SARS-CoV-2 variants and sarbecoviruses^7,8^. These findings suggest that vaccines targeting the conserved S2 region can induce immunity against future variants and other emerging coronaviruses.

**Figure 1.**
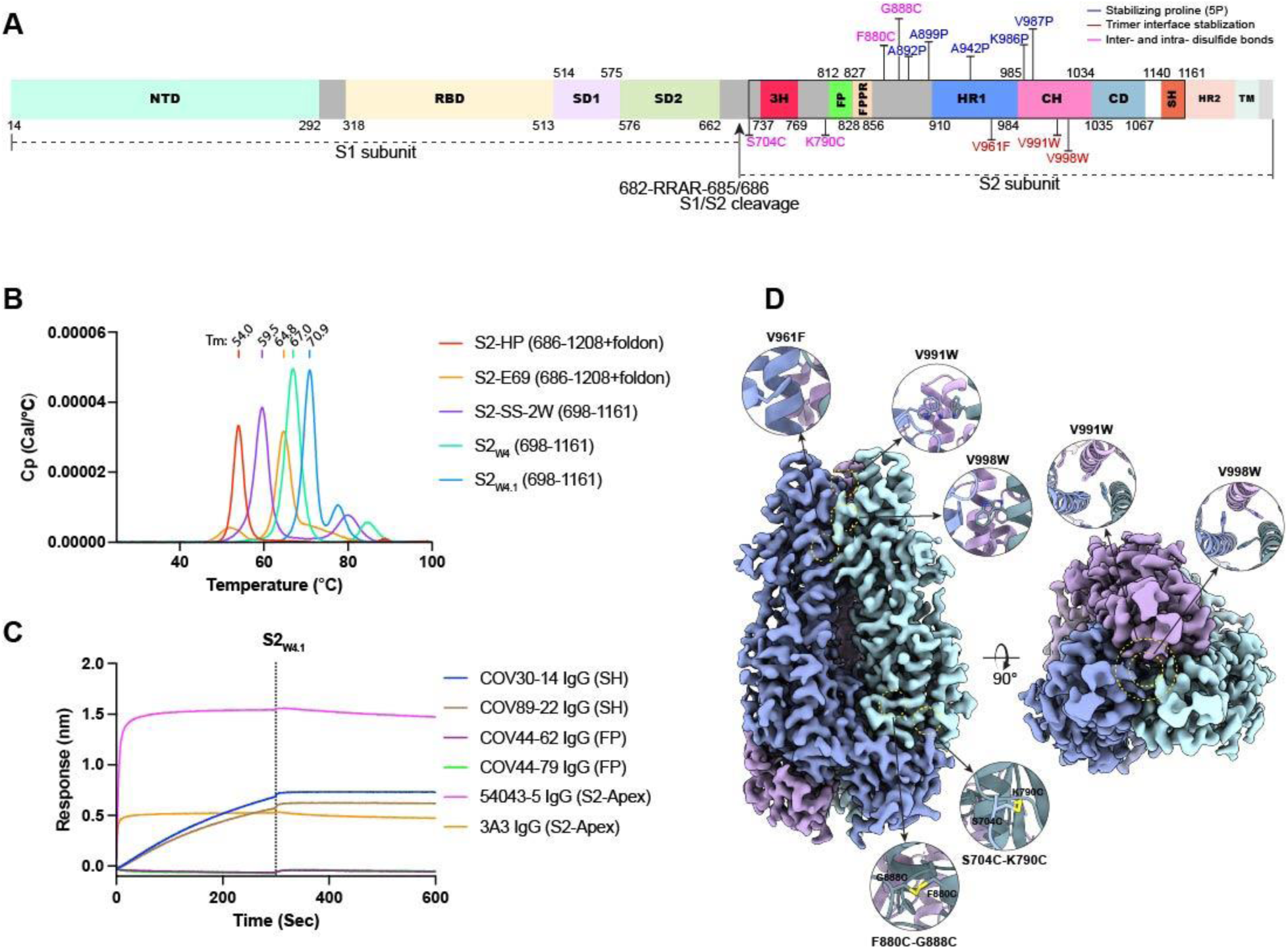
Design and characterization of stabilized S2_W4.1_ construct. (A) Schematic diagram of SARS-CoV-2 spike S2 subunit stabilization. The diagram highlights key stabilizing mutations introduced to enhance the stability of the S2 subunit. Each substitution is annotated with a colored label denoting its functional category as shown in the upper right legend. Spike domain regions are labeled: NTD (N-terminal domain), RBD (receptor-binding domain), SD1/SD2 (subdomain 1/2), 3H (3-helix bundle), FP (fusion peptide), FPPR (fusion peptide proximal region), HR1 (heptad repeat 1), CH (central helix), CD (connector domain), SH (stem helix), HR2 (heptad repeat 2), and TM (transmembrane domain). The S1/S2 furin cleavage site (682-RRAR-685/686) separates the S1 and S2 subunits. (B) Differential scanning calorimetry profiles of stabilized S2 proteins. S2_W4.1_ is compared with other stabilized S2 constructs, including previously reported stabilized S2. The substitution lists of all stabilized S2 are provided in Table S1. (C) Biolayer interferometry (BLI) analysis of S2_W4.1_ binding to SARS-CoV-2 S2-targeting antibodies. Antibodies are labeled by their binding epitopes (SH, FP, and S2-Apex regions). (D) Cryo-EM structure of S2_W4.1_ protein shown in two orientations. The left panel displays the side view, where the viral membrane if present would be at the bottom, and the right panel shows the top view down the trimer axis of the same structure with the three subunits in different colors (cyan, blue, purple). Mutations introduced for S2 stabilization are highlighted in cartoon representation with side chains shown as sticks, and their positions on S2 are marked by yellow dashed circles. For V991W and V998W, both side and top views are shown.

The major epitopes identified in S2 to date can be categorized into three regions: the stem helix (SH), the fusion peptide (FP), and the apex of S2 (S2-Apex). Numerous broadly neutralizing antibodies targeting the SH and FP regions have been isolated^9–16^. While their neutralization potency is not as high as S1 antibodies, they demonstrate pan-neutralizing activity sufficient to neutralize across, or even beyond, the betacoronavirus genus. In contrast, the S2-Apex region has been relatively poorly characterized, with only a few antibodies recently reported to bind this region^17–22^. However, these S2-Apex binding antibodies exhibit high antigen reactivity, sufficient to bind broadly spike proteins from multiple members of the *Orthocoronaviridae*, but demonstrate minimal neutralizing activity, resulting in limited protective effects.

The spike protein is metastable, which is a feature of viral fusion proteins^23^. The S2 subunit is structurally unstable in the prefusion state and tends to rapidly rearrange into an elongated postfusion conformation when the S1 domain is removed, similar to the influenza virus hemagglutinin (HA) stem^24–27^ and HIV gp41^28–31^. Therefore, stabilizing the S2 subunit to maintain its prefusion structure is essential for studying S2 antigenicity or utilizing it as a vaccine. To overcome this challenge, antigens with prefusion S2 stabilization mutations have been developed^19,26,32–34^. Although these stabilized S2 proteins have facilitated the investigation of the conserved S2 subunit, our understanding of polyclonal-level immune responses to respective S2 epitopes induced by infection and vaccination remains limited.

Our previously reported EMPEM technology^35–37^ has enabled polyclonal antibody (pAb) epitope mapping against a variety of microbial antigens at the three-dimensional structural level, and this technique has continued to advance^38–40^. Moreover, the recent integration of ModelAngelo (MA) technology^41^ with structure-to-sequence (STS) analysis^39^ has made it possible to deduce protein sequences from high-resolution structures obtained through cryo-EMPEM^42,43^.

Here, we generated a stabilized S2 protein through the fusion of previously characterized stabilization residues and employed it to perform epitope mapping of pAbs to S2 using negative-stain EMPEM (ns-EMPEM) with plasma samples from SARS-CoV-2-infected and vaccinated individuals. Furthermore, we deduced the sequences of monoclonal antibodies (mAbs) targeting the immunodominant epitope through STS analysis using cryo-EMPEM. These results, together with antibody database and B-cell receptor (BCR) sequence analyses, demonstrate that the S2-Apex is a highly immunodominant region targeted by antibodies derived mainly from the IGHV3-30 family, which is one of the most frequently utilized germlines among coronavirus antibodies^44–48^. These antibodies, derived from the IGHV3-30 germline, also converge on acquiring a specific CDRH3 motif and represent one of the most dominant antibody clonotypes elicited in coronavirus vaccine recipients.

## RESULTS

### Stabilization of SARS-CoV-2 S2 domain

To investigate the immune response against the SARS-CoV-2 S2 domain, we designed a stabilized prefusion S2 protein. Prior research has reported various approaches for stabilizing the SARS-CoV-2 S2 domain^19,26,33,34^. In this study, we engineered an S2 domain construct with improved stability by combining previously reported mutations with additional stabilizing residues (hereafter referred to as S2_W4.1_) **(Figure 1A and Table S1)**. This construct, encompassing residues 698-1161 of the spike protein, incorporated five proline mutations from the traditionally used hexa-proline stabilization: A892P, A899P, A942P, K986P, and V987P^49–51^. Additionally, three substitutions were introduced at the inter-protomeric interface, T961F, V991W, and T998W, for expression of an intact (closed) trimeric S2 domain ^33,34^, and an inter-subunit disulfide bond was engineered via S704C and K790C substitutions to prevent trimer dissociation^26^. To further enhance protomeric stability and folding, G880C and F888C mutations were incorporated to create an intra-subunit disulfide bond. The S2_W4.1_ expressed as a stable trimer despite eliminating the C-terminal T4 fibritin motif (foldon), which is conventionally used to stabilize trimerization of the spike protein to resemble its native trimeric form. The thermal stability of S2_W4.1_ protein was estimated by differential scanning calorimetry (DSC). In comparison with other stabilized S2 spike proteins (S2-SS-2W, S2-E69)^33,34^ and Hexapro S2 (S2-HP) with melting temperatures (*T*_m_) of 59.5, 64.8, and 54.0°C, respectively, S2_W4.1_ showed improved thermal stability, displaying a high *T*_m_ value of 70.9°C **(Figure 1B)**. In the absence of V991W and T998W (S2_W4_), the construct exhibited a *T*_m_ of 67.0°C.

To further evaluate the antigenic integrity of S2_W4.1_, binding against known S2 binding antibodies was assessed using a bio-layer interferometry (BLI) assay. When tested for binding to mAbs targeting S2-Apex, FP, and SH (S2-Apex: 54043-5, 3A3; FP: COV44-62, COV44-79; SH: COV89-22, COV30-14)^9,13,18,21^, S2_W4.1_ retained binding to S2-Apex and SH-binding antibodies, while losing binding to FP antibodies **(Figure 1C)**. This impaired antibody binding to the FP could be caused by the proline substitutions incorporated for spike stabilization, as previously described by Dacon *et al.*^13^ Indeed, binding to FP antibodies was rescued when K986P and V987P mutations (2P) were reverted back to wild-type residues **(Figure S1)**. In the absence of 2P stabilizing mutations, the S2 apex region becomes unstable or more flexible, potentially leading to exposure of the FP region. Consequently, structural studies assessing S2-reactive antibodies were carried out using S2_W4.1_, supplemented with FP peptide-based binding studies to assess antibody reactivity to FP.

To further resolve the structural features of S2_W4.1_, we determined the apo S2 structure using cryo-EM at a resolution of 2.75 Å **(Figure 1D and Table S2)**. Structural comparison with other SARS-CoV-2 S2 domain prefusion structures (PDB ID: 8VQB and 8VAO) revealed a root-mean-squared deviation (Cα RMSD) of 0.94 Å and 0.90 Å, respectively **(Figure S2A)**. The overall structures exhibited high similarity, except in the inherently dynamic areas. Notably, structural comparison of S2_W4.1_ with the S2 domain of full-length spike (PDB ID 6XR8) revealed conformational differences in residues 828-856 (FPPR, fusion-peptide proximal region) **(Figure S2A and B)**. In the absence of S1, the FPPR tends to preferentially adopt a downward conformation that is incompatible with the S1 domain due to steric hindrance. This conformational change can also be observed in previously reported spike S2-only structures^19,26,33,34^. These results thus provide structural and functional validation of our S2_W4.1_ protein for subsequent EM-based mapping of antibody responses to the SARS-CoV-2 S2 domain.

### The apex region of SARS-CoV-2 S2 represents the most immunodominant S2 epitope

Plasma from nine adult study participants (referred to as Donors 1-9) with varying SARS-CoV-2 immune histories was analyzed, including individuals with immune encounters limited to COVID-19 immunizations (V, vaccine), limited to documented SARS-CoV-2 infection (I, infection) or both (VI). . Additionally, longitudinal plasma from the same participants collected at different time points were used to study changes in their antigenic profiles over time **(Figure 2A and Table S3)**. To map the overall immune response to S2 in these donors, ns-EMPEM was performed with S2_W4.1_ complexed with Fabs derived from purified polyclonal IgGs from each plasma sample. Interestingly, S2 antibody reactivity across all donors using S2_W4.1_ as a probe was primarily focused on the apex region, although they possessed varying antibody titers against the spike protein **(Figure 2B and Table S4)**. Further analysis of ns-EMPEM results revealed that antibody responses to the S2-Apex targeted two distinct sites. Antibodies to the first site bind in a nearly vertical orientation to the apex of the S2 domain (red), while antibodies to the second site approach from a more lateral direction (blue) (**Figure 2B**). We designated these as the Apex-A and Apex-B sites, respectively. Antibody responses to both sites were observed across all samples, regardless of whether they were derived from the V group (Donors 1, 2, and 3), I group (Donors 6 and 7) or both (Donors 4, 5, 8, 9) **(Figure 2A and B)**. These antibodies were also detected in biospecimens collected at the early stage (10 days post-infection) and persisted for more than 539 days following the last vaccination (Donor 6 and Donor 1, respectively), indicating that they are induced during the initial phase and maintained as part of the long-term immune response. In the ns-EMPEM study, all detectable antibody responses were focused on the S2-Apex, with no reactivity at alternative epitopes. These results are consistent with immunodominant responses to the S2-Apex (FP-binding epitope could not be evaluated in this analysis and will be discussed below).

**Figure 2.**
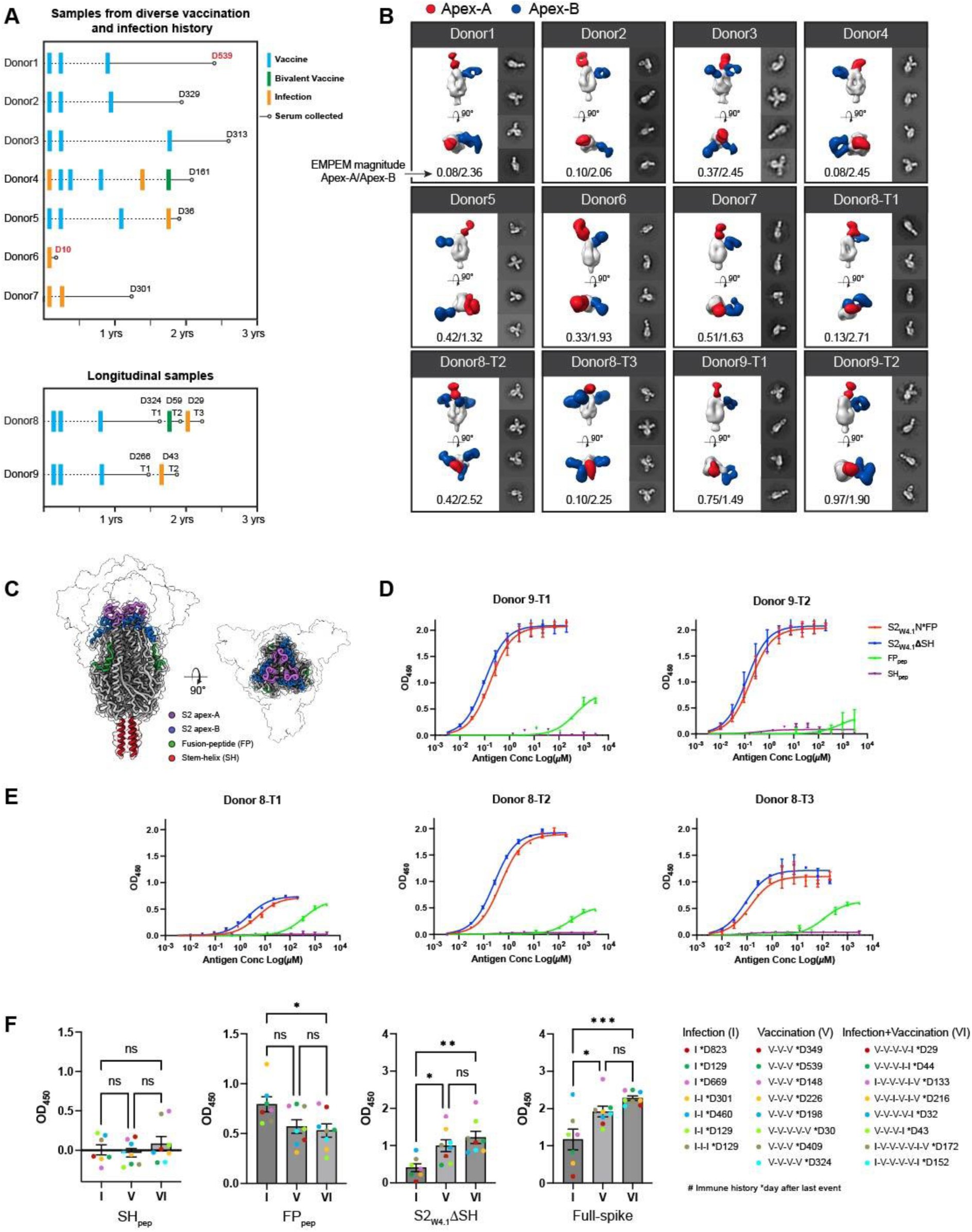
Ns-EMPEM analysis and immunodominance of the S2-Apex. (A) Timeline of plasma collection from nine donors, annotated with vaccination, bivalent vaccine booster, and infection events. For each donor, the day of plasma collection is labeled as the number of days after the most recent immune event. The shortest (D10) and longest (D539) intervals after the last event are highlighted in red. Longitudinal sampling points (T1-T3) are indicated for donor 8 and donor 9. (B) Ns-EMPEM reconstructions of polyclonal antibody responses mapped onto the prefusion S2 trimer. Representative side and top views of the 3D composite map and 2D classes are displayed in a square from each donor sample. In the 3D map, the S2 protein is colored gray, and the Apex-A and Apex-B binding antibodies are colored red and blue, respectively. The EMPEM magnitude of the Apex-A and Apex-B epitopes for each dataset is listed and described in detail in the Methods section. (C) Schematic representation of S2 domain antibody epitopes. Distinct epitope regions are indicated by different-colored cartoon models, and the full-length spike, including S1, is shown as a white surface model. (D-E) ELISA analysis of plasma antibody titers against individual S2 epitopes. Longitudinal plasma samples are analyzed for donor 9 (D) and donor 8 (E). Different probes were used to assess epitope-specific responses: S2_W4.1_N*FP for total S2-binding antibodies, S2_W4.1_𝛥SH for apex-directed antibodies, FP_pep_ for fusion peptide antibodies, and SH_pep_ for stem helix antibodies. Data are shown as mean ± SD from two independent experiments (each performed in duplicate). (F) Single-point ELISA measuring plasma antibody binding to SH_pep_, FP_pep_, S2_W4.1_𝛥SH, and full-length spike to assess relative antibody levels at each epitope. Each dot represents an individual donor sample (n = 23 donors; I = 7, V = 8, VI = 8). Bars indicate mean OD_450_ with SEM. Donors are grouped as infection only (I), vaccination only (V), or vaccination plus infection (VI). The donor key shown in the rightmost panel specifies the immune history and the number of days after the last immune event when plasma was collected, with colors corresponding to individual samples in the graphs. Statistical significance is determined by one-way ANOVA with Tukey’s multiple comparisons test (* *p* < 0.05, ** *p* < 0.01, *** *p* < 0.001; ns, not significant).

The EMPEM magnitude (average number of Fabs per trimer) for the Apex-A and Apex-B epitopes reveals differential binding occupancy **(Figures 2B and S3A)**. Apex-A shows an average of 0.36 Fabs per trimer, while Apex-B demonstrates substantially higher occupancy with 2.09 Fabs per trime**r**. In 2D and 3D classifications, Apex-A binding antibodies were not observed with more than one Fab bound per trimer, whereas Apex-B binding antibodies showed up to three Fabs simultaneously engaged on a single prefusion S2 trimer. However, this restricted binding stoichiometry may not indicate a low frequency of Apex-A directed antibodies but rather reflects a limitation in their simultaneous engagement within a single prefusion trimer. The observation that Apex-A antibodies bind to only a single protomer per trimer suggests that simultaneous engagement of two or more protomers within the same trimer may destabilize S2, rendering such complexes undetectable. In addition, 3D classification reveals that Fab densities at the Apex-A epitope exhibit considerable angular variation **(Figure S3B)**. This heterogeneity may reflect either the diversity of Apex-A-binding antibodies in the polyclonal response or inherent flexibility of the epitope itself. In contrast, Apex-B-binding antibodies displayed homogeneous Fab binding densities, indicating that this epitope preferentially accommodates antibodies with a specific binding angle or approach geometry **(Figure S3C)**.

The stabilized S2_W4.1_ probe used for epitope mapping does not exhibit reactivity to antibodies targeting the FP region **(Figure 1C)**. To assess whether the S2-Apex region is, indeed, immunodominant over other S2 epitopes, we evaluated the presence of FP-binding and other antibodies in these samples by enzyme-linked immunosorbent assay (ELISA). The epitopes identified to date within the S2 domain include S2-Apex, FP, and the SH region **(Figure 2C)**. To quantify relative antibody titers to these epitopes, we generated various probes to effectively distinguish responses to each of these sites **(Figure S4A)**. Given that S2_W4.1_ protein already lacks binding capacity for FP-targeting antibodies, further removal of S2_W4.1_ SH region enables it to be specific for detecting S2-Apex binding antibodies. To achieve this, S2_W4.1_ΔSH was generated by truncating residues 1141-1161, which constitute the SH region. To comprehensively detect all antibodies binding to the S2 domain, we further designed the S2_W4.1_N*FP construct, in which the FP epitope region (residues 812-827) was incorporated at the N-terminus to allow for efficient detection of FP-targeting antibodies. In addition, synthetic peptides including the SH (residues 1138-1167) and FP (residues 808-837) epitope regions were employed to identify antibodies specific for each respective site. Since the probes were constructed in different formats, we assessed their ability to reliability quantify antibody specificities by testing them with equal amounts of their corresponding monoclonal antibodies. Increasing the amount of each antigen probe demonstrated that the antibody response reached comparable Bmax values. While the binding kinetics may vary depending on the probe or antibodies, the Bmax values indicate that relative antibody quantities can be reliably assessed **(Figure S4B)**.

In addition, binding of antibodies specific to each S2 epitope (FP, SH, and S2-Apex) was evaluated by ELISA using the S2_W4.1_ probe **(Figure S4C)**. The FP-targeting antibody showed no detectable binding to S2_W4.1_, indicating that FP binding is impaired in this construct. In contrast, the SH-binding antibody exhibited clear binding, and ns-EM analysis further confirmed that SH-targeting antibodies can be visualized and identified by ns-EM **(Figure S4D)**. Using the designed probes, antibody responses to the S2 epitopes were evaluated in longitudinal plasma samples from Donor 8 and Donor 9 by ELISA **(Figure 2D and E)**. Overall, antibodies against the SH and FP regions were substantially lower compared to those against the S2-Apex. The low SH and FP responses were further validated by the similar binding response in plasma reactivity to S2_W4.1_ΔSH as compared to S2_W4. 1_N*FP. In congruence with our EMPEM results, responses targeting the S2-Apex region were highly dominant compared to other sites. Donor 8 sera showed an especially dramatic increase in apex-directed antibodies following additional vaccination (bivalent) after T1, whereas responses to FP and SH remained unchanged. Similarly, while Donor 9 sera exhibited minimal changes in the overall antibody responses between T1 and T2 even after natural infection between these timepoints (**Figure 2A**), a robust response against S2-Apex was observed across both time points. Collectively, these findings demonstrate the immunodominance of S2-Apex over other S2 sites and indicate that despite being buried under the S1 domain in the full spike, the S2-Apex is sufficiently accessible to elicit a strong antibody response.

To examine how antibody responses varied relative to different SARS-CoV-2 immune events, binding titers against each epitope were assessed in the I, V, and VI groups using the ELISA **(Figure 2F)**. In this analysis, we compared the binding profiles of seven sera from group I and eight sera each from groups VI and V across four antigen constructs: FP, SH, S2-Apex, and full-length spike. Analysis revealed that antibodies against the SH region showed no major differences among groups, whereas FP-targeting antibodies exhibited significantly higher titers in the I group than in the VI group. These results demonstrate that the 2P stabilization of spike employed by most SARS-CoV-2 vaccines presents the fusion peptide inefficiently, leading to differences in antibody responses between the I and V/VI groups. In contrast, antibody titers against the S2-Apex region were substantially increased in the V and VI groups compared with the I group. This increase followed the same pattern as the overall antibody response induced by full-length spike. These findings demonstrate that multiple vaccinations and the combination of infection and vaccination are more effective at eliciting S2-Apex-targeting antibodies than infection alone.

### Cryo-EMPEM analysis revealed that antibodies targeting the S2 Apex-B epitope are derived from the IGHV3-30 germline gene

To further characterize these S2-Apex antibodies, we employed a structure-guided sequence analysis approach using cryo-EMPEM with plasma-derived pAbs from Donor 1, Donor 2, Donor 3, and Donor 8-T2. Apex-A antibodies were detected in samples from Donor 3 and Donor 8-T2, and the corresponding structures were determined at 3.25 Å and 4.24 Å resolution, respectively **(Figures 3A, S5C, and S5F)**. Furthermore, high-resolution structures of S2_W4.1_ in complex with Apex-B binding pAbs were resolved at 3.17, 3.16, 3.21, and 2.83 Å resolution from Donor 1, Donor 2, Donor 3, and Donor 8-T2, respectively **(Figure 3B, S5A, S5B, S5E, S5G, and Table S2)**. Although Apex-A antibodies were also detected in Donor 1 and Donor 2 sera by ns-EMPEM, no corresponding structures were obtained in the cryo-EMPEM analysis. As discussed above, this can be explained by the relatively low binding magnitude of Apex-A antibodies observed in Donor 1 and Donor 2 by ns-EMPEM, as well as by either competitive binding of multiple antibodies at the Apex-A site or conformational flexibility, which may have further limited map resolvability. Likewise, in the Apex-A binding antibody structures, the local resolution at the epitope-paratope interface was lower than that observed for the Apex-B antibodies **(Figures S5C, D, and F)**. Even in the Donor 3 dataset, where the overall map was resolved to 3.25 Å, the Fab density was not sufficiently clear to allow atomic model building. Upon additional 3D classification of the S2 Apex-A pAb complex from Donor 3, we resolved two distinct classes of Apex-A binding antibody engaging the same epitope **(Figures S5C, D, and H)**. For both classes, the relatively low resolution of the antibody region hampered atomic model interpretation for the Fab. Superimposition of the two EM maps revealed no detectable structural changes in the S2 apex, but the approach angle of antibody binding differed by approximately 17° when measured between the center of mass of each Fab and C𝛼 of Asp985 (centering Apex A epitope). In contrast, in the S2 Apex-B structures, poly-alanine models of the Fab regions were built for each cryo-EMPEM map **(Figure 3C)**. When superimposed, these models showed highly similar approach angles and overlapping binding epitopes, indicating convergent recognition of the S2-Apex across donors.

**Figure 3.**
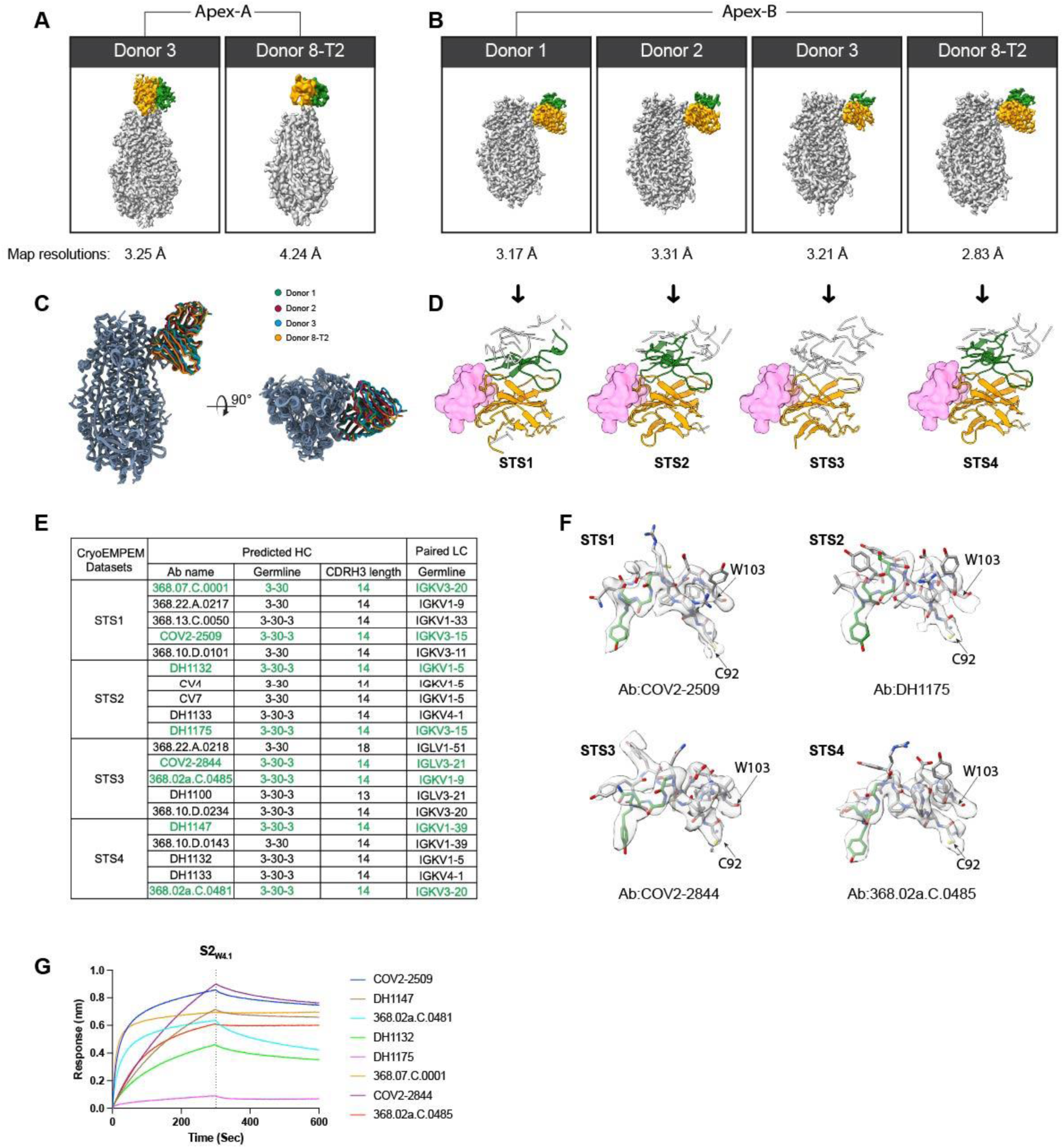
Cryo-EMPEM analysis and structure-based sequence prediction of Apex-B antibodies. (A-B) Focused cryo-EMPEM refinement analysis of S2 protein-antibody complexes targeting distinct epitope regions. (A) Apex-A epitope-specific antibodies from Donor 3 and Donor 8-T2. (B) Apex-B epitope-specific antibodies from Donors 1, 2, 3, and 8-T2. Resolutions determined by Fourier shell correlation are indicated for each reconstruction (Figure S4). The S2 protein is shown in gray with bound antibody densities highlighted in orange (HC) and green (LC). (C) Similar binding modes of Apex-B targeting antibodies. Overlay of four S2-Fab structures derived from the maps in panel (B), shown in both side view and top view orientations. The Fab regions were built as alanine models and are color-coded according to donor: Donor 1 (green), Donor 2 (red), Donor 3 (blue), and Donor 8-T2 (orange). (D) 3D models of the antibody parts from each cryo-EMPEM dataset built by ModelAngelo (MA). Only the predicted regions containing secondary structure, as determined by MA, are highlighted in yellow for the HC and in green for the LC, while fragmented regions are shown in gray. The corresponding epitope regions are displayed as pink surface models. Each dataset used in the sequence-to-structure (STS) analysis is marked as STS1, STS2, STS3, and STS4. (E) Predicted antibody list from STS analysis. Expressed antibodies for further study are colored in green. (F) The CDRH3 density maps from the STS1, 2, 3, and 4 datasets are shown as transparent gray surfaces. Antibody sequences predicted from STS analysis were used for model building for CDRH3 (STS1: COV2-2509; STS2: DH1175; STS3: COV2-2844; STS4: 368.02a.c.04; see Figure 3D). The G/S-G-S/N-Y motif is highlighted in green in the stick model (related to Figure 4). The positions of flanking residues C92 and W103 of CDRH3 are labeled in each structure. (G) BLI analysis of predicted antibodies against the S2_W4.1_ protein. Eight IgG antibodies were analyzed with S2_W4.1_, and the corresponding sensorgrams are shown in different colors for each antibody.

Next, we used the high-resolution structural information from EM densities of Apex-B antibodies to predict their sequences. This approach combined the previously reported structure-to-sequence (STS) analysis with ModelAngelo (MA), enabling a more integrated framework for sequence inference directly from structural data^39,41,42^ **(Figure 3B and D**; STS1, STS2, STS3, and STS4). First, MA, an AI-based model builder, was used to predict the sequence based on the map densities. The no_seq option of MA allowed successful prediction of the entire pAb heavy chain regions, while the light chain predictions generated severely fragmented models due to relatively lower resolutions of this region. This could either reflect the flexibility of the light chains caused by their limited contribution in S2 engagement or due to their sequence diversity. For each heavy chain, MA generated fragmented structures and corresponding sequences as hidden Markov models (HMM) files. The fragmented structures were reordered by comparison with a known Fab structure, and the HMM files were subsequently integrated following this arrangement to produce a unified sequence that was visualized **(Figure S6A).** The integrated HMMs were matched to sequences from the coronavirus antibody database (CoV-AbDab)^52^ using the HMMER search function in MA. From the results, five antibodies with the lowest E-value scores were selected for each dataset **(Figure 3E)**, none of which had their epitopes defined in the original data. Sequence alignment of the database antibodies with the predicted sequences revealed an average sequence identity of approximately 50% **(Figure S6B)**. Remarkably, all matched antibodies were encoded by the IGHV3-30 germline gene family, and all sequences except for two displayed an identical CDRH3 length of 14 amino acid residues **(Figure 3E)**. In contrast, the paired light chain germlines were diverse, underscoring the pronounced heavy chain germline restriction in antibody recognition of the Apex-B epitope. When one of the selected antibodies was built into the CDRH3 density maps of each STS dataset, the amino acid sequence showed substantial agreement with both the length and the positional alignment of the CDRH3 residues in the density map **(Figure 3F)**. Clear densities were observed for the N-terminal Cys92 and the C-terminal Trp103 flanking the CDRH3, within which 14 residues were accurately built.

To determine whether these identified antibodies could engage the Apex-B epitope and to also investigate possible light-chain preference, we expressed and purified eight antibodies in total, selecting two clones from each dataset employing distinct light chains (from STS1: 368.07.C.0001 and COV2-2509; from STS2: DH1132 and DH1175; from STS3: COV2-2844 and 368.02a.C.0485; from STS4: DH1147 and 368.02a.C.0481)^53–55^. Notably, all expressed antibodies demonstrated binding to the S2 protein by BLI assay, although binding to DH1175 was much lower than the others **(Figure 3G).** Subsequently, ns-EM and BLI-based competition assays were performed to confirm their epitope specificities. Ns-EM analysis demonstrated that both COV2-2509 and COV2-2844 engaged the Apex-B binding site, while the remaining antibodies were indirectly shown to target the same epitope through BLI-based competitive assays **(Figures S6C and D)**. Collectively, cryo-EMPEM structure-guided sequence prediction with MA enabled the identification of eight antibodies that bound to a previously uncharacterized epitope, all of which were found to share the IGHV3-30 family germline.

### IGHV3-30 antibodies with 14-residue CDRH3s exhibit sequence convergence and robust clonal expansion targeting Apex-B

Examination of germline usage frequency within the CoV-AbDab shows that IGHV3-30 family is the most represented (16.2%), which aligns with prior literature indicating that IGHV3-30 constitutes one of the most prevalent antibodies in the repertoire of SARS-CoV-2-infected individuals **(Figure 4A)**^44–48,56^. While most of these studies have emphasized the importance of IGHV3-30 family antibodies as neutralizing antibodies targeting the S1 domain,^45,46,57^ we have shown that the highly immunodominant S2-Apex response also utilizes the IGHV3-30 germline and specifically employs 14-residue-long CDRH3s, which closely approximates the reported average CDRH3 length of 14.8 residues in the naive human antibody repertoire^58^. To further analyze the features of the IGHV3-30 in CoV-related antibody repertoires, the IGHV3-30 family antibodies in the database were initially classified based on their specificity to S1, S2, or unknown categories using epitope annotations from CoV-AbDab, with S1 representing the largest proportion at 52%, followed by unknown at 29% and S2 at 19%. Strikingly, when the analysis was restricted to antibodies with a CDRH3 length of 14 residues, the majority shifted to S2 and unknown categories at 37% and 50%, respectively, with S1 reduced to 13% **(Figure 4B)**. Sequence alignment of CDRH3 regions from the eight Apex-B binding antibodies (368.07.C.0001, COV2-2509, DH1132, DH1175, COV2-2844, 368.02a.C.0485, DH1147, and 368.02a.C.0481) revealed a common G/S-G-S/N-Y motif at positions 5-8 of CDRH3. In particular, the glycine at position six and tyrosine at position eight were identical in all antibodies **(Figure 4C)**. To examine the preference of tyrosine at position 8 of CDRH3, we filtered for antibodies with a 14-residue-long CDRH3 with a tyrosine at the 8th position. The database antibodies showed that S2 and unknown categories accounted for 41% and 55%, respectively, while S1 was markedly reduced to 4%. Additionally, since the G and Y residues at positions 6 and 8 of the CDRH3 are simultaneously conserved within the G/S-G-S/N-Y motif described above, we further applied a filter for the GxY pattern. Under this criterion, S2 and unknown categories accounted for 41% and 56%, respectively, while S1 slightly decreased to 3%. This result is largely consistent with the previous filtering by the 8th-position tyrosine alone, indicating that residues G at position 6 and Y at position 8 are highly co-conserved. Subsequently, to examine the CDRH3 sequence conservation of 540 antibodies commonly using the IGHV3-30 family with 14-residue CDRH3, corresponding to the second column of Figure 4B, these antibodies were grouped by epitope and aligned **(Figures 4D-F)**. While no discernible commonalities were observed among the 70 antibodies targeting S1, the 199 S2-targeting antibodies displayed the same G/S-G-S/N-Y motif as observed in Apex-B binding antibodies **(Figures 4D and E)**. Interestingly, this conserved sequence pattern was also maintained among the unknown epitope antibodies, suggesting that many of these antibodies are likely enriched for Apex-B binding specificities **(Figure 4F)**. Moreover, the number of antibodies with 14-residue CDRH3 comprises approximately 30% of all IGHV3-30 family antibodies in the COVID-19 mAb database (CoV-AbDab), constituting the largest subset, of which over 80% encode tyrosine at the 8th residue **(Figure 4G)**. These observations suggest that IGHV3-30 antibodies exhibit strong CDRH3 sequence convergence. This pattern of CDRH3 conservation and S2 binding preference in IGHV3-30 genes, including IGHV3-30/IGHD1-26 combinations, is consistent with our previous report^47^. To determine whether Apex-B binding antibodies exhibit a particular IGHD gene preference, we examined the DNA sequences of the eight identified antibodies. Here we show that the antibodies utilized a variety of IGHD gene combinations, including IGHD1-26, IGHD2-15, IGHD3-10, and IGHD3-16, with no apparent preference. Especially, in cases where the tyrosine residue at position 8 of the CDRH3 was not contributed by the IGHD gene (COV2-2844, 368.02a.C.0485, and DH1147), this residue was compensated by the long IGHJ6 gene **(Figure S7A)**. Therefore, the characteristic 14-residue-long CDRH3 with the G/S-G-S/N-Y motif observed in Apex-B binding antibodies can be generated through IGHV3-30 in combination with various D and J genes, which likely underlie the ease with which these antibodies are elicited and their dominance in the repertoire.

**Figure 4.**
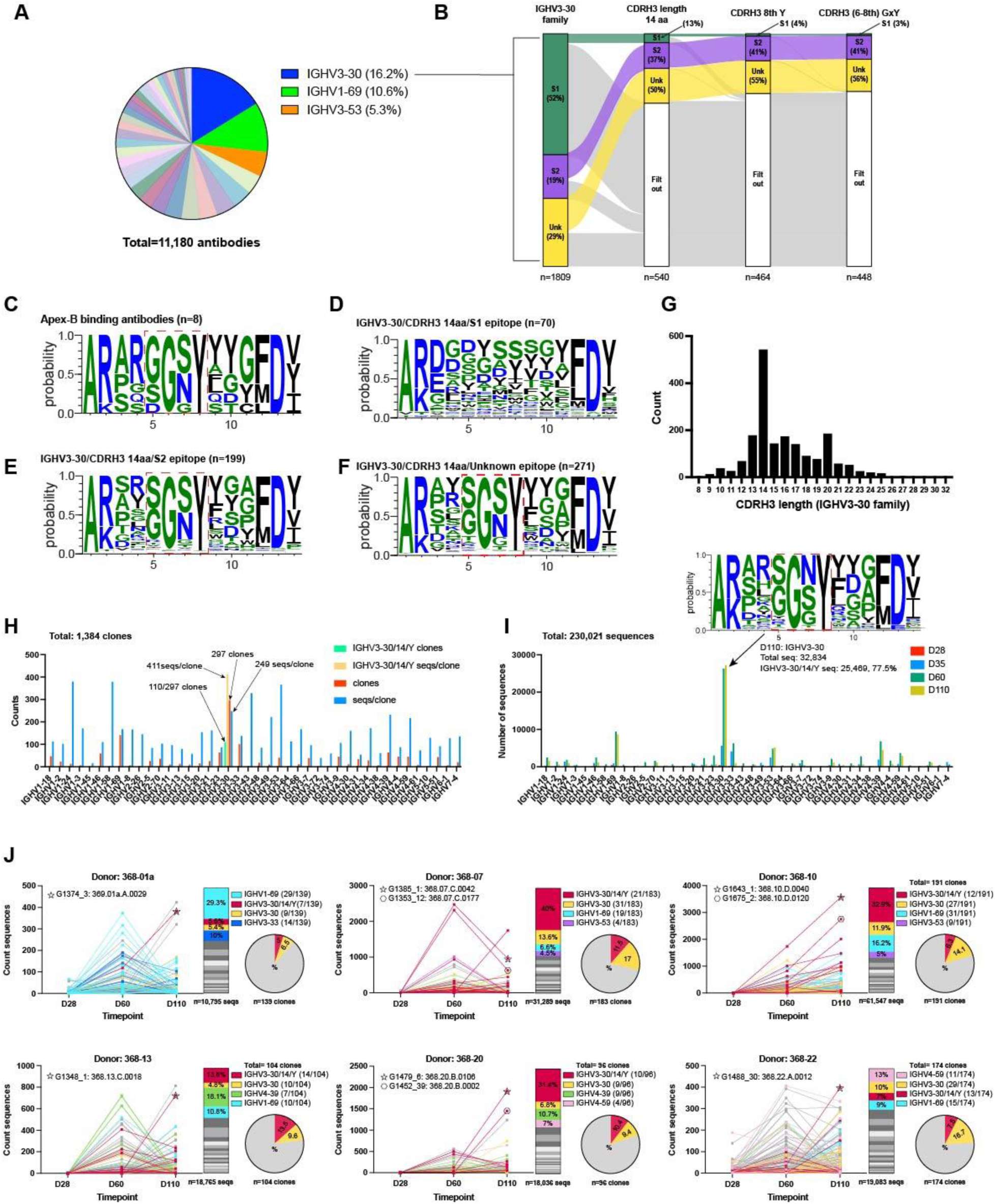
CoV-AbDab and BCR sequence analysis of IGHV3-30-biased Apex-B antibodies. (A-G) Antibody sequence analysis using CoV-AbDab. (A) Pie chart showing IGHV gene distribution of 11,180 antibody sequences from CoV-AbDab. (B) Alluvial plot showing the epitope distribution of IGHV3-30 subgroup antibodies under successive filtering conditions. The analysis begins with the initial IGHV3-30 subgroup distribution, followed by filtering for CDRH3 length of 14 amino acids, and further filtering for tyrosine at the 8th position of CDRH3, and for glycine and tyrosine at the 6th and 8th positions (GxY motif). The n values below each column indicate the number of antibodies remaining after filtering, and numbers in parentheses denote percentages for each epitope antibody. Unk = unknown; Filt = filtered. (C) Sequence logo plot of the CDRH3 regions from eight Apex-B binding antibodies identified in Figure 3E. The conserved G/S-G-S/N-Y motif is highlighted with a red dashed box. (D-F) Sequence logo plots of IGHV3-30 antibodies with a CDRH3 length of 14 amino acids, classified by epitope specificity. S1 epitope (D), unknown epitope (E), and S2 epitope (F) subsets correspond to the CDRH3 length 14 aa groups shown in (B). The conserved G/S-G-S/N-Y motif is highlighted with a red box. (G) Distribution of CDRH3 lengths among IGHV3-30 subgroup antibodies in the CoV-AbDab. (H-J) BCR sequencing analysis results using data from Kim et al.^53^. (H) Analysis of unique clone counts and sequences per clone for each IGHV germline across eight different donors. Clone counts are shown in red and sequences per clone in blue. IGHV3-30/14/Y-featured clones are highlighted in green and orange, with specific annotations. IGHV3-30 clone counts include IGHV3-30/14/Y clones. (I) Analysis of sequence expansion over time across different IGHV gene families from eight donors. Bar chart showing the expansion of each IGHV gene across different time points (Day 28, Day 35, Day 60, and Day 110). The accompanying sequence logo displays the consensus motif derived from 25,469 sequences of IGHV3-30/14/Y at D110. (J) Longitudinal tracking of clonal expansion in six individual donors (368-01a, 368-07, 368-10, 368-13, 368-20, and 368-22) across three post-vaccination time points (D28, D60, and D110). Each dot represents a unique clone at each time point, with clones from the same clonal lineage connected by lines across time points. For each donor, the top IGHV germline clones with high sequence abundance are displayed in different colors, while remaining clones are shown in gray. Pie charts display the proportional distribution of IGHV3-30/14/Y-featured and IGHV3-30 clones among total clone numbers. IGHV3-30 represents non-IGHV3-30/14/Y clones only. Antibodies expressed for epitope characterization are labeled as clone ID: antibody name and marked at D110 with star and hexagon symbols. n = total sequences or clones.

Further analysis of the eight expressed antibodies revealed that DH1147, DH1132, DH1175, 368.02a.C.0481, and 368.02a.C.0485 shared CDRH1 and CDRH2 sequences identical to the IGHV3-30 germline, with DH1132 and DH1175 showing 100% identity across all of the IGHV gene coding region **(Figure S7B)**. Collectively, these antibodies can recognize the epitope in a naive germline configuration with a CDRH3 of 14 amino acids, consistent with the typical repertoire average length, and accommodate diverse light-chain pairings, further supporting their ready propensity for expansion within the antibody repertoire.

Through CoV-AbDab data analysis, we found that antibody sequences with the IGHV3-30 family, having CDRH3 consisting of 14-length amino acids and tyrosine at the 8th residue, converge into the Apex-B binding G/S-G-S/N-Y motif (IGHV3-30/14/Y). Antibodies classified by these characteristics were obtained from 27 independent references, with the largest contribution of 211 antibodies reported by Kim *et al.* **(Table S5)**^53^. To assess whether this convergence also occurs within individual repertoires, we further analyzed the sequencing dataset from Kim *et al.* The BCR sequencing data were derived from eight donors who received prime and boost vaccinations at 0 and 3 weeks, respectively. First, all clones across the eight donor repertoires were analyzed, yielding a total of 1,384 unique clones **(Figure 4H)**. Among these, the IGHV3-30 family germline accounted for 297 clones, representing the largest clone number compared to other germlines. Within IGHV3-30, 110 clones possessed the IGHV3-30/14/Y characteristic, corresponding to 37% of all IGHV3-30 clones. Although the average number of sequences per clone was not markedly higher for the IGHV3-30 family relative to other germlines, the IGHV3-30/14/Y subset stood out, with an average of 441 sequences per clone. This analysis indicates that the IGHV3-30/14/Y clones drive the overall clonal expansion within the IGHV3-30 family and, moreover, display the highest degree of clonal expansion across all IGHV germlines analyzed. Subsequently, we examined sequence expansion over time for each IGHV germline **(Figure 4I)**. Strikingly, IGHV3-30 sequences showed substantial expansion after day 35. By day 110, 77.5% of all IGHV3-30 sequences were found to carry the IGHV3-30/14/Y characteristic, demonstrating that the clonal expansion observed within IGHV3-30 was largely driven by these clones. Additionally, their CDRH3 regions displayed a highly conserved G/S-G-S/N-Y motif, with the G and Y residues co-conserved in 99% of all IGHV3-30/14-featured sequences. This observation is consistent with the above CoV-AbDab analysis.

Furthermore, analysis of IGHD and IGHJ gene usage among IGHV3-30 sequences carrying the 14-residue CDRH3 and the G/S-G-S/N-Y motif characteristic of Apex-B-binding antibodies revealed a broad diversity of D-J gene combinations **(Figure S7C)**. All IGHJ genes except IGHJ2 were used, and multiple D genes including IGHD1-7, IGHD1-26, IGHD2-15, IGHD2-21, IGHD3-10, IGHD3-16, IGHD3-22, IGHD4-17, IGHD4-23, IGHD5-12, and IGHD6-6 were compatible with this feature, with IGHD1-26 being the most frequently used. This observation further supports the notion that the dominance of Apex-B targeting antibodies is facilitated by a broad range of D-J gene combinations rather than being restricted to specific germline pairings.

Using these BCR sequencing datasets, we further assessed the clonal expansion and prevalence of clones having the IGHV3-30/14/Y feature from six individual donors **(Figure 4J)**. Across multiple donors, clonal expansion was predominantly driven by IGHV3-30 family clones. Notably, IGHV3-30 family clones comprised 53.6%, 44.8%, 18.6%, and 38.2% of total repertoire sequences in individual donors 368-07, 368-10, 368-13, and 368-20, respectively.

Within this, IGHV3-30/14/Y-featured clones were particularly enriched, comprising 40.0%, 32.9%, 13.8%, and 31.4% of the total sequences in the same donors, highlighting their striking expansion. In addition, these clones exhibited marked sequence expansion at day 60 and remained dominant through day 110, indicating their long-term immune activity. At the clone level, these IGHV3-30/14/Y clones accounted for 21 of 183 (11.5%), 12 of 191 (6.3%), 14 of 104 (13.5%), and 10 of 96 (10.4%) unique clones in the respective donors. In donors 368-01a and 368-22, overall expansions of IGHV1-69 and IGHV4-59 clones, respectively, were more pronounced, yet a single IGHV3-30/14/Y clone continued to exhibit the highest clonal expansion at day 110. To confirm that the expanded IGHV3-30/14/Y clones specifically recognize the Apex-B epitope, we selected the most highly expanded clones from each donor **(Figures 4J and S8)**. In total, nine representative antibodies were expressed and purified, and their binding activity to S2_W4.1_ was evaluated using the BLI assay. All nine antibodies showed clear binding to S2_W4.1_ **(Figure S8A)**. Moreover, competition assays with the COV2-2509 antibody demonstrated that each antibody competes for the same binding site, indicating that they all target the Apex-B epitope **(Figure S8B)**.

In addition, we compared IGHV3-30/14/Y-encoded B-cell receptors across healthy, infected, and vaccinated donors using BCR sequencing data from Briney *et al*. and the Observed Antibody Space (OAS) database^59,60^. When compared with precursor frequencies reported for other antiviral antibodies, IGHV3-30/14/Y BCRs exhibited a relatively high baseline frequency in pre-pandemic healthy donors, with a median of 56.2 clones per million BCRs^61,62^ **(Figure S8C)**. Following SARS-CoV-2 infection or vaccination, the frequency of these clones increased substantially, reaching 517.3 and 1,593.8 clones per million BCRs, respectively. Notably, vaccination was associated with an approximately threefold higher frequency than infection, indicating that vaccination may support the presence of a larger pool of IGHV3-30/14/Y clones. Importantly, these values reflect increases in unique clonotypes independent of clonal expansion; when considering donor-specific clonal expansion patterns observed in our analyses, the total abundance of these antibodies is expected to be substantially higher.

Together, these findings indicate that IGHV3-30, particularly the IGHV3-30/14/Y subset, undergoes preferential selection and expansion through antigen-driven processes that favor this germline usage across multiple individuals. Given that these sequencing data were obtained from vaccinated individuals, these results further suggest that vaccination promotes robust expansion of Apex-B binding antibodies, representing a convergent and durable immune response centered on the Apex-B epitope.

### Structural characteristics of S2 Apex-B epitope-antibody interactions

To investigate how antibodies targeting the Apex-B epitope utilize the IGHV3-30 germline sequence for binding and to assess the importance of the characteristic IGHV3-30/14/Y feature, we determined the cryo-EM structure of the COV2-2509 mAb in complex with S2_W4.1_ **(Figures 5A and Table S2)**. The COV2-2509 antibody recognizes a discontinuous conformational epitope in the S2 domain of the spike protein, spanning residues 737-765 encircling the outer apex region and residues 845-853. CDRH1 and CDRH2 recognize the 737-765 region, while CDRH3 bridges the 737-765 and 845-853 segments together with CDRL3. The HC contributes to a buried surface area (BSA) of 766 Å², compared with only 103 Å² from the LC, indicating that the interaction is predominantly mediated by the HC. This HC-dominant binding provides a rationale for the ability of Apex-B antibodies to utilize diverse light chain pairings. The interface is stabilized by hydrogen bonds, salt bridges, and hydrophobic contacts, largely mediated by CDRH2 and CDRH3 **(Figure 5A-C)**. Key residues include Tyr52A, Asp53, Asp55, Tyr58, Gly97, Tyr100, and Gln100A (CDRH1:26-33; CDRH2:51-54; CDRH3:93-102), which participate in hydrogen bonding, while Asp55 and Lys64 contribute to salt bridge formation. In contrast, for the light chain, Trp94 of CDHL3 is the sole residue involved in hydrophobic interactions. The major hydrophobic contacts, established by HC Val50, Tyr58, and Tyr100 together with LC Trp94, form a hydrophobic pocket that encloses S2 Leu849, where the aromatic ring surfaces of Tyr residues and Trp are oriented toward the interior of the pocket, creating a nonpolar environment and establishing hydrophobic interactions that stabilize the interface. However, the involvement of Trp94 in binding does not appear to be essential. In the COV2-2509 antibody, the tryptophan residue is located within CDRL3 and encoded by the IGKV3-15 gene. In contrast, Apex-B binding antibodies can arise from diverse IGKV/IGLV combinations, and the presence of tryptophan at this position is not conserved **(Figure S9A)**. Moreover, the light chains of the 17 distinct Apex-B binding antibodies identified in this study exhibit variable CDRL3 lengths ranging from 7 to 12 residues. These observations indicate that Apex-B binding antibodies are not restricted to a particular light chain but can be formed through a variety of light-chain combinations.

**Figure 5.**
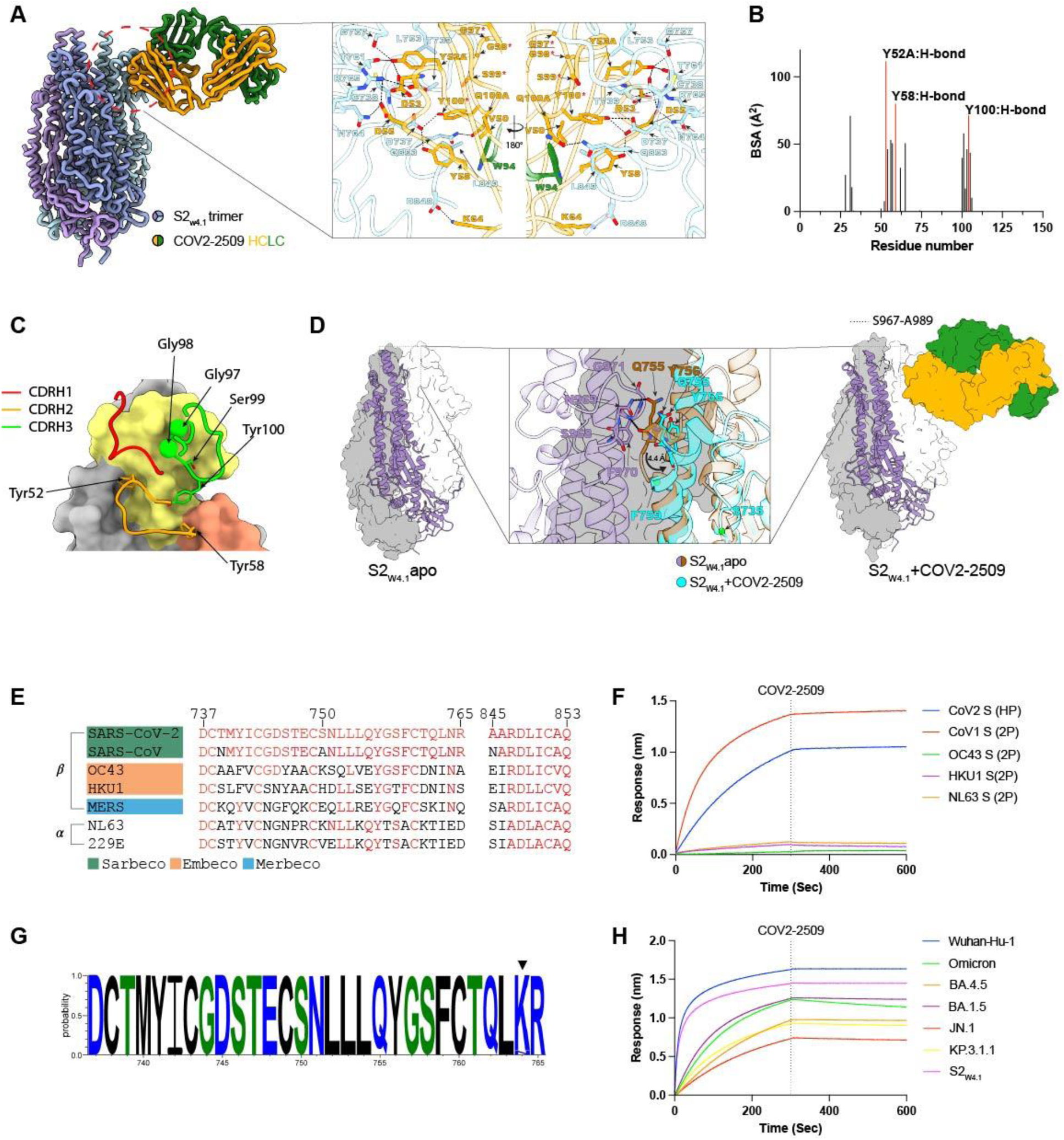
Cryo-EM structure of S2_W4.1_ in complex with COV2-2509 Fab and its binding analysis. (A) Side view of the atomic model built into the 3.1 Å cryo-EM map shown as a cartoon representation with the membrane proximal region at the bottom. The binding interface between S2 and Fab is enlarged in two views, with key interacting residues displayed as sticks. Hydrogen bonds and salt bridges are indicated by dashed lines, and residues belonging to the G/S-G-S/N-Y motif are marked with red asterisks and labeled with residue numbers. Main-chain interaction residues are underlined below their residue numbers. (B) Graph showing the buried surface area (BSA) contributions of HC residues at the Fab-S2 interface. The three tyrosine residues with the highest BSA values are highlighted in red, and each participates in hydrogen bonding. (C) CDRH1, CDRH2, and CDRH3 are shown as cartoon models, while the S2 region is represented as a surface model. The 737-765 epitope region is colored yellow, and the 845-853 region in salmon. Three key tyrosine residues and residues of the G/S-G-S/N-Y motif are shown as sticks, with glycines represented as spheres. (D) Structural comparison of S2_W4.1_ apo and S2_W4.1_-COV2-2509 complexes. Left, S2_W4.1_ apo structure; Light, S2_W4.1_-COV2-2509 complex structure. The COV2-2509 Fab is depicted in yellow and green surface model, and the missing residues S967-A989 are indicated by a dashed line. Center, zoomed-in view of the boxed region with the antibody-bound protomer (cyan) superimposed on the apo structure (brown). Key residues are shown as sticks, hydrogen bonds as dashed lines, and S735 and F759 as green spheres. (E) Sequence alignment of the COV2-2509 antibody binding region across human coronaviruses (hCoVs). Residues conserved relative to the SARS-CoV-2 sequence are highlighted in red. *Alphacoronaviruses* and *betacoronaviruses* are labeled as α and β, respectively. Among *betacoronaviruses*, the Sarbecovirus, Embecovirus, and Merbecovirus lineages are color-coded in green, orange, and blue, respectively. (F) BLI binding analysis of COV2-2509 IgG against spike proteins from hCoVs, including SARS-CoV-2 (CoV2 S), SARS-CoV-1 (CoV1 S), OC43, HKU1, and NL63. (G) Sequence logo plot of 56 SARS-CoV-2 variants. The Asn764Lys substitution is indicated with a triangle above the corresponding residue. (H) BLI binding analysis of COV2-2509 IgG with spike proteins from six SARS-CoV-2 variants, including Wuhan-Hu-1 as a reference.

Notably, Tyr52A in CDRH2, Tyr58 in HFR3, and Tyr100 in CDRH3 establish extensive hydrogen bonds and hydrophobic interactions, through their hydroxyl groups and aromatic rings, respectively, contributing substantially to the binding interface. These three residues also account for the largest BSA of the HC **(Figure 5B and C)**. Importantly, Tyr52A and Tyr58 of CDRH2 are encoded by the IGHV3-30 germline, and Tyr100 of CDRH3 was conserved across all Apex-B binding antibodies found in this study. To assess whether the two tyrosine residues, Tyr52A and Tyr58, are unique features of the IGHV3-30 family, we aligned and compared all 467 unmutated human IGHV germline reference sequences, including all known alleles **(Figure S9B)**^63^. The analysis revealed that Tyr58, located in HFR3, is highly conserved across nearly all IGHV germlines. In contrast, Tyr52A is conserved only within the IGHV3-30, IGHV3-30-3, IGHV3-30-5, and IGHV3-33 genes **(Figure S9C)**. Notably, although both Tyr52A and Tyr58 are present in these four genes, IGHV3-33 differs by possessing a bulky tryptophan residue at position 52 (instead of Ser52), immediately preceding Tyr52A, which likely introduces steric hindrance that prevents optimal engagement of the Apex-B epitope **(Figure S9D)**. Similarly, the IGHV3-30*02 and IGHV3-30-5*02 alleles contain arginine instead of Ser52, which also causes steric hindrance with surrounding residues due to its long side chain. Collectively, the combination of Tyr52A and Tyr58 residues critical for Apex-B binding is specifically preserved in IGHV3-30, IGHV3-30-3, and IGHV3-30-5 germlines, excluding the IGHV3-30*02 and IGHV3-30-5*02 alleles.

Gly97, Gly98, and Ser99 within the G/S-G-S/N-Y motif play a key role in establishing structural complementarity of CDRH3 with the S2 domain, together with the characteristic 14-residue CDRH3 observed in Apex-B binding antibodies **(Figure 5C)**. Notably, Gly98 is strictly conserved across all Apex-B binding antibodies, and substitution with residues that carry larger side chains than glycine would potentially result in steric interference with CDRH1, leading to alterations in CDRH3 conformation. In particular, Gly98 adopts an unusual region of the Ramachandran plot with a strongly positive phi angle of +123.3 degrees (Figure S9E). This conformation is strongly restricted because non-glycine residues would introduce steric clashes between the C beta atom and the backbone carbonyl groups. Moreover, Gly98 is completely buried with a solvent accessible surface area of 0.0 Å^2^, placing it within the structural core of the beta-turn. Together, this buried position and glycine-specific backbone geometry explain why Gly102 is critically conserved within the G/S-G-S/N-Y motif.

Next, we compared the structures of S2_W4.1_ alone and in complex with COV2-2509 **(Figure 5D)**. This analysis revealed that COV2-2509 binding to the S2 Apex-B epitope induces structural rearrangements within Ser735-Phe759 **(Figure 5D)**. In addition, in the COV2-2509-bound structure, the S967-A989 loop of the neighboring protomer is disordered. In the apo structure, this region is stabilized through side-chain interactions between N969 and Q755 of the adjacent protomer, as well as main-chain interactions between the backbone amide of F970 and the carbonyl of Q755. Further stabilization arises from van der Waals contacts between S968-G971 and Q755-Y756 of the adjacent protomer. Upon COV2-2509 binding, however, Q755 undergoes a displacement of approximately 4.4 Å (measured at the Cα), and the Ser735-Phe759 segment shifts outward. These conformational changes are anticipated to destabilize the S967-A989 region. Additionally, when the full-length spike (CoV2-HP-S) was incubated with COV2-2509 and analyzed by ns-EM, the spike trimer was observed to dissociate into monomers **(Figure S10A-C)**. This dissociation could be the result of destabilization of the Ser735-Phe759 region, which could act as a molecular latch for maintaining trimeric integrity.

Within the Apex-B binding site of the S2 domain, residues 845-853 are highly conserved across all human coronaviruses, including SARS-CoV-2, human coronaviruses OC43, HKU1, NL63, and 229E, and Middle East respiratory syndrome coronavirus (MERS-CoV). In contrast, the 737-765 region exhibits sequence divergence across alpha-CoVs (NL63, 229E) and non-sarbecovirus beta-CoVs, including *Embecovirus* (OC43, HKU1) and *Merbecovirus* (MERS-CoV), but remains highly conserved among *Sarbecoviruses* (SARS-CoV-1, GZ02, WIV1, Rp3, HKU3-1, RaTG13, ZXC32, and YN2018C [Pangolin-CoV-GD]) **(Figures 5E and S10D)**. BLI experiments show that the COV2-2509 antibody binds strongly to SARS-CoV-1 and SARS-CoV-2 but does not bind to OC43, HKU1, or NL63 seasonal coronaviruses **(Figure 5F)**. Moreover, alignment of more than 50 SARS-CoV-2 variants, including recent lineages, showed that all residues within the 737-756 region are conserved, except for the Asn764Lys substitution found in Omicron and subsequent lineages **(Figure 5G)**. BLI results demonstrated that the COV2-2509 antibody maintained high binding across all spike proteins tested, from the ancestral strain through the recent KP3.1.1 variant **(Figure 5H)**. Taken together, these results reveal that three tyrosine residues (Tyr52A, Tyr58, and Tyr100) with the G/S-G-S/N-Y motif are important in mediating interaction with the Apex-B epitope. In addition, these antibodies were experimentally confirmed to bind SARS-CoV-1 and SARS-CoV-2, and the conservation of their binding site across sarbecoviruses and SARS-CoV-2 subvariants suggests a broad potential for cross-sarbecovirus recognition.

### Preferential Apex-B binding of COV2-2509 to cleaved spikes

The Apex-B epitope is buried within the S1 domain in the context of the full-length spike. Structural overlay of the S2_W4.1_-COV2-2509 Fab complex structure onto the full-length spike structure (PDB ID 6XR8) revealed that the COV2-2509 epitope is located at the apex of the S2 domain, hidden inside the region between the NTD and the RBD of the S1 domain **(Figure 6A)**. This indicates that Apex-B binding antibodies cannot access their epitope when the S1 domain remains in place. To investigate whether COV2-2509 binding is promoted by S1/S2 cleavage and S1 dissociation, we employed several spike constructs. These included spike HP variants containing either the native S1/S2 cleavage site sequence RRAR (S-HP-RRAR; RRAR residues 682-685) or a GSAS substitution (S-HP-GSAS), which blocks furin-mediated cleavage. In addition, to prevent trimer dissociation, we used spike HP mut7 constructs^64^ containing an inter-protomer disulfide bond (S-HPM7-RRAR and S-HPM7-GSAS). We first evaluated the structural diversity of the cleaved spike ectodomain alone (S-HP-RRAR) by ns-EM analysis as a reference. Interestingly, in a subset of particles, we observed an entirely open S1 conformation in which one S1 protomer was displaced outward, thereby providing the possibility of exposing the otherwise hidden Apex-B epitope **(Figure 6B)**. While representing only a small subset of particles, these observations suggest that conformational breathing of the S1 domain could provide transient access to the buried Apex-B epitope.

**Figure 6.**
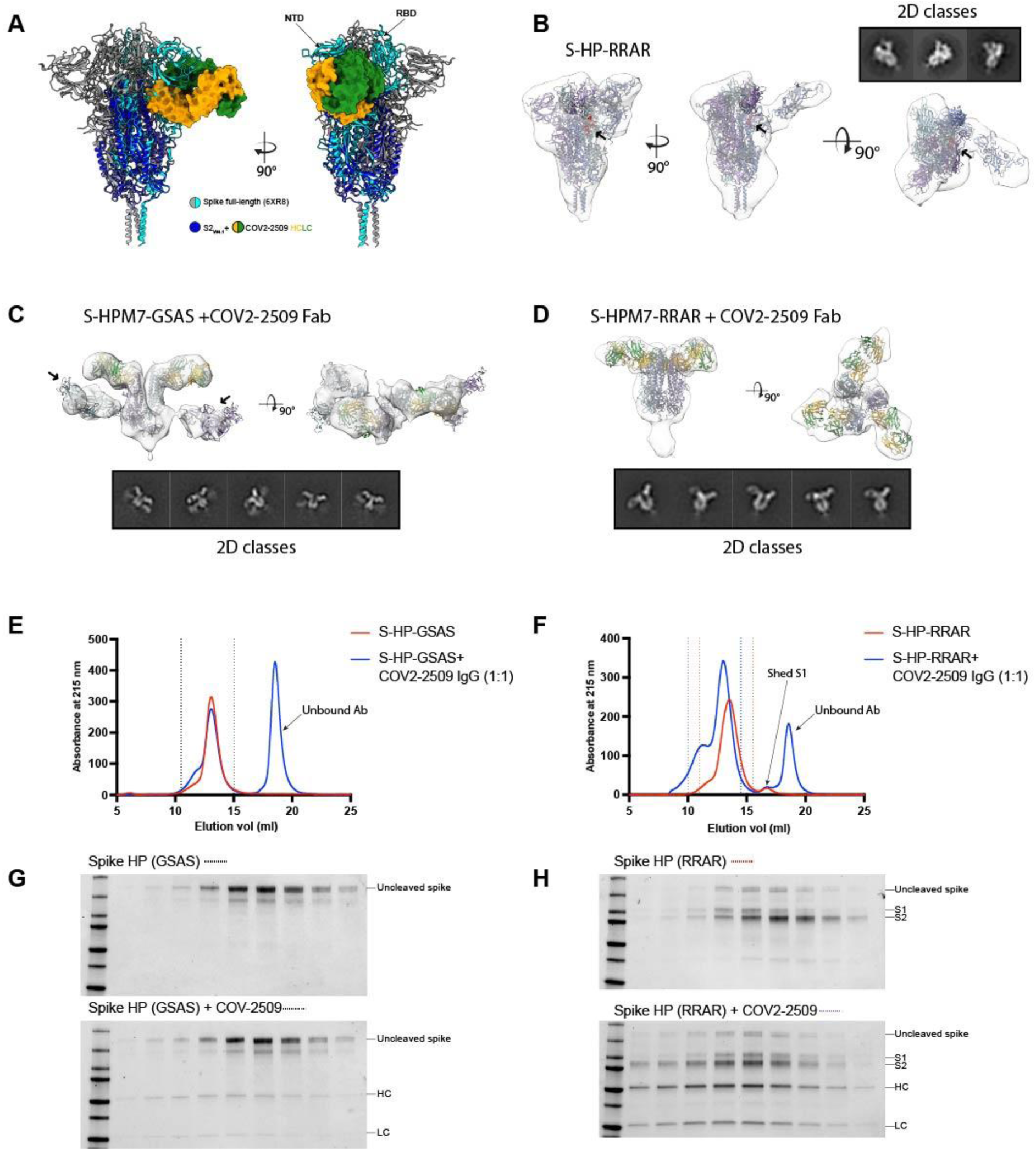
Recognition of the Apex-B epitope by COV2-2509 in the full-length spike. (A) Structural comparison of the S2_W4.1_-COV2-2509 complex with the full-length spike (PDB ID 6XR8). The S2_W4.1_ trimer is shown in blue, with the COV2-2509 Fab depicted as surface representation in yellow and green for HC and LC, respectively. In the full-length spike trimer, one protomer is highlighted in cyan and the remaining protomers are shown in gray. The NTD and RBD of the cyan protomer are indicated by arrows. (B-D) Representative 2D classes and ns-EM reconstructions of S-HP-RRAR alone (B), S-HPM7-GSAS or S-HPM7-RRAR in a complex with COV2-2509 Fab (C, D). Modeled atomic structures were fitted into the 3D reconstruction for visualization. In (B), the Apex-B epitope is highlighted in red with an arrow, and in (C), the S1 domain is indicated by arrows. (E, F) Size-exclusion chromatograph (SEC) profiles of S-HP (GSAS or RRAR) in the absence or presence of COV2-2509. Peaks corresponding to unbound antibody and shed S1 are indicated. The elution volume of the fractions collected for SDS-PAGE analysis is marked with dashed lines. (G, H) SDS-PAGE of SEC fractions from (E, F), with the corresponding dashed lines and colors indicated above each gel image.

Next, we examined COV2-2509 binding in relation to S1/S2 cleavage using ns-EM. Ns-EM analysis showed that, upon binding of COV2-2509 Fab to S-HPM7-GSAS, the S1 domain densities were observed flanking the S2 domain, indicating that the S1 domain was displaced from its native position **(Figure 6C)**. In contrast, no S1 density was detected in the S-HPM7-RRAR-COV2-2509 Fab complex, indicating complete release of S1 following cleavage **(Figure 6D)**. These observations support the notion that displacement of S1 from its native position is required to allow antibody access to the Apex-B epitope. However, despite increasing the antibody concentration to more than a threefold molar ratio over spike and extending incubation for over 24 hours, a substantial fraction of unbound spike particles remained.

We further assessed binding in solution by size exclusion chromatography (SEC) using the original HP spike (S-HP-RRAR), which does not contain inter-protomer disulfide bonds. We analyzed both S-HP-GSAS and the cleaved S-HP-RRAR variants. In the S-HP-GSAS construct, most antibodies failed to form complexes, whereas in the cleaved S-HP-RRAR construct, nearly all spikes shifted to the antibody-spike complex peak **(Figures 6E-H)**. This finding indicates that S1/S2 cleavage facilitates exposure of the Apex-B region. Notably, additional S1 shedding was not observed upon antibody binding, suggesting that COV2-2509 does not actively induce S1 dissociation **(Figure 6F and H)**. Instead, the antibody appears to preferentially engage spike species that have already undergone partial S1 shedding, which are enriched in the cleaved S-HP-RRAR population. Supporting this notion, the sodium dodecyl sulfate-polyacrylamide gel electrophoresis (SDS-PAGE) result of S-HP-RRAR apo showed that S1 and S2 were present at unequal ratios, with disproportionately stronger S2 bands in the later SEC fractions, indicating a relative loss of S1 **(Figure 6H)**. Additionally, the persistence of S1 despite antibody binding is expected to arise from antibody engagement with only one or two protomers of partially shed spike trimers.

Therefore, Apex-B binding antibodies are expected to preferentially engage partially shed spikes, in which S1 has been shed from one or more protomers within the trimer, a form that may exist in sufficient numbers to permit expansion of Apex-B responses once S1/S2 cleavage has occurred. Moreover, these antibodies are also likely to bind a subset of spikes that undergo outward opening of the S1 domain through breathing motions^21,65,66^, as supported by recent MD simulation results showing that S1/S2 cleavage promotes large-scale S1 movement and expansion of the S2 central helix^67^, and by prior reports of spontaneous S1 shedding in native spikes^25,68^. Therefore, although the spike protein is stabilized in the prefusion state, the presence of a fraction of spikes in a partially shed state could induce excessive amplification of Apex-B antibodies, as noted above.

### Absence of neutralizing function of S2 Apex-B binding antibodies

We next determined the neutralization and Fc-mediated properties of seven S2 Apex-B binding antibodies (COV2-2509, COV2-2844, DH1132, DH1147, 368.02.a.C.0481, 368.07.C.0001, 368.02a.C.0485). In parallel, we assessed two additional control S2-directed antibodies targeting distinct regions, COV44-79 (FP-mAb) and COV89-22 (SH-mAb)^9,13^. None of the Apex-B binding antibodies tested exhibited measurable entry inhibition of SARS-CoV-1 or SARS-CoV-2 pseudoviruses **(Figure 7A and B)**, nor did they show microneutralization activity against replication-competent ancestral SARS-CoV-2 virus **(Figure 7C)**. In our studies, the control mAb COV44-79 exhibited neutralizing activity against SARS-CoV-2 pseudovirus entry (EC₅₀ = 1.85 µg/mL), whereas COV89-22 showed low neutralizing activity against the SARS-CoV-1 pseudovirus (EC₅₀ = 11.42 µg/mL) **(Figure 7A and B)**. Notably, neither antibody exhibited measurable inhibition of live virus in our microneutralization assay **(Figure 7C)**. These results highlight the low neutralization potency of S2-binding antibodies. However, all seven Apex-B binding antibodies exhibit strong antibody-dependent cellular cytotoxicity (ADCC) activity **(Figure 7D)**. Thus, while the Apex-B antibodies fail to neutralize viral entry and replication, their robust antibody-dependent cellular cytotoxicity points to efficient engagement of Fcγ receptors. These properties are consistent with features previously reported for multiple S2-Apex binding antibodies^17–20,22^. Such antibodies generally exhibit little to no neutralizing and protection activity but display high ADCC activity. Therefore, S2 Apex-B binding antibodies are non-neutralizing, yet may retain limited protective potential through Fc-mediated effector functions.

**Figure 7.**
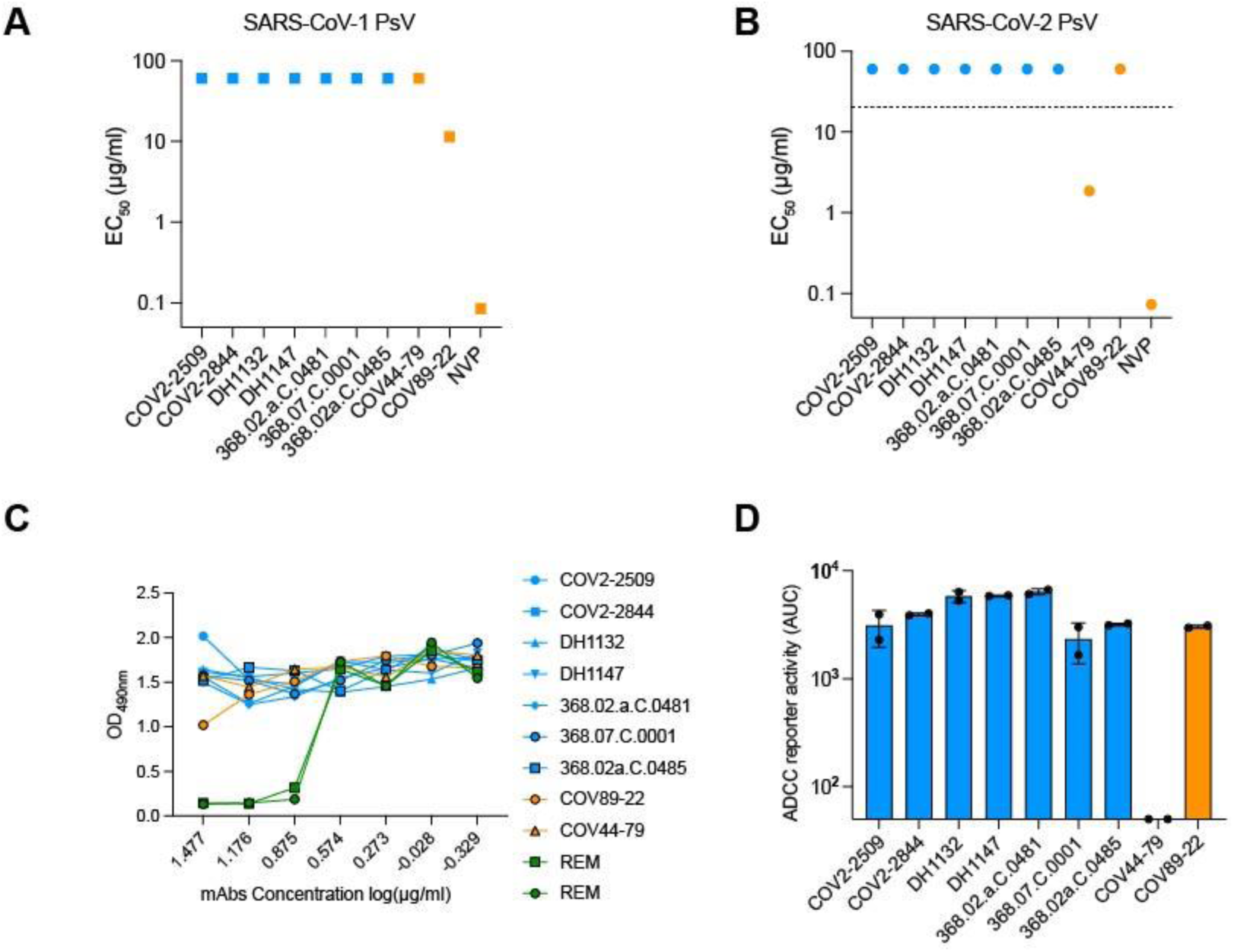
Neutralization and Fc-mediated effector functions of S2 Apex-B monoclonal antibodies. (A-B) Neutralization activity of S2 Apex-B binding antibodies against SARS-CoV-1 (A) and ancestral SARS-CoV-2 (B) spike-pseudotyped viruses. EC₅₀ values are shown for each antibody, and the dashed line indicates the assay detection limit. Across all panels, Apex-B binding antibodies are shown in blue, FP (COV44-79) and SH (COV89-22) direct antibodies are shown in orange, and the positive controls nevirapine (NVP) and remdesivir (REM) are shown in green. The dashed line denotes background activity. Data are shown as mean ± s.d. (C) Live SARS-CoV-2 multicycle microneutralization assay showing dose-dependent inhibition of viral infection by S2 Apex-B antibodies, measured as nucleoprotein signal (OD₄₉₀). (D) Antibody-dependent cellular cytotoxicity (ADCC) activity of S2 monoclonal antibodies, expressed as area under the curve (AUC).

## DISCUSSION

The continuous evolution of coronaviruses underscores the need for vaccines that can provide broad and durable protection. While the strategy of repeatedly updating vaccines to match emerging variants can provide some level of protection, a precision vaccine targeting conserved antigenic sites that do not drift would be transformative. Therefore, precision vaccine design should prioritize conserved antigenic sites that are more likely to elicit protective antiviral immune responses. In this regard, the S2 subunit of the spike protein exhibits high sequence conservation across *Orthocoronaviridae* and has emerged as a key target for the development of next-generation universal coronavirus vaccines. Achieving this objective requires a deeper understanding of S2-specific immune responses.

This study analyzed sera from donors with diverse immune histories in a structure-based manner and identified the previously underappreciated S2-Apex as the most immunodominant target within the S2 subunit. Although it is masked by S1 in the prefusion conformation, it is likely transiently accessible through S1 breathing motions^65,69^ or partial shedding, thereby exposing the immunodominant epitope. These findings demonstrate that structurally occluded antigenic sites can nonetheless serve as potent focal points for antibody responses. Moreover, we demonstrate that antibodies targeting Apex-B are found to converge into a public antibody class characterized by IGHV3-30/14/Y features with a conserved 14-residue CDRH3 G/S-G-S/N-Y motif, and they are repeatedly observed across multiple donors by antibody databases and BCR sequencing analyses. The reasons these antibodies exhibit remarkable immunodominance can be summarized in four points: (i) they possess strong binding capacity even in their germline-encoded state, eliminating the need for substantial somatic hypermutation (SHM); (ii) they tolerate diverse light-chain pairings, enabling multiple B-cell clones to readily participate in this response; (iii) conserved CDRH3 motifs can be repeatedly generated through diverse VDJ recombination pathways; (iv) finally, consistent with our observation that IGHV3-30/14/Y-encoded B-cell receptors are already present at appreciable frequencies (median 56.2 clones per million BCRs; Figure S8C) in pre-pandemic healthy donors, previous reports have shown that IGHV3-30 is intrinsically predisposed to preferential utilization during V(D)J recombination^70^, positioning these germlines for rapid amplification upon antigen exposure. The combination of these properties underlies why IGHV3-30/14/Y-featured antibodies are found to be public antibodies to SARS-CoV-2 and Sarbecoviruses and display remarkable expansion capability in targeting Apex-B. As a result, this clonotype undergoes robust expansion in multiple donors and emerges as a highly dominant component of the antibody repertoire. In line with these explanations, previous analyses of B-cell repertoires from convalescent COVID-19 patients have revealed that IGHV3-30 family germlines, including IGHV3-30-3, are utilized at substantially higher frequencies than other germline genes, and these frequencies markedly exceed those observed in healthy controls^44,45,47,48^, in agreement with our findings. A similar phenomenon is also observed for IGHV3-53/3-66-class antibodies directed against the SARS-CoV-2 RBD, which repeatedly appear in different individuals as a prototypical public antibody class and exhibit high immunodominance among RBD-targeting antibodies^56^. However, IGHV3-30 family clones recognizing the Apex-B epitope represent the most immunodominant S2-directed antibody population among coronavirus vaccine recipients. In some individuals, these clones can account for up to 40% of total spike-reactive sequences, surpassing even the most dominant S1-directed responses such as IGHV3-53/3-66 RBD antibodies (which represent ∼10% of total RBD-binding antibodies). This suggests that the Apex-B epitope is a critical region that must be carefully considered in future vaccine design.

Across multiple independent studies, B-cell responses to SARS-CoV-2 infection or vaccination consistently converge on the S2 region and predominantly elicit non-neutralizing antibodies. Nevertheless, the immunological significance of this recurring S2-focused response has remained underexplored, largely because prior research emphasized the S1 subunit, which is dominant for neutralizing antibody responses, as the primary correlate of protection. In Sakharkar *et al*., neutralizing antibody levels declined over time^71^. In contrast, S2-specific memory B cells were maintained or expanded and were dominated by non-neutralizing antibodies of the IGHV3-30 lineage. Sequence analysis using the IgG memory B cells (MBCs) sequencing dataset of Sakharkar *et al*. revealed that 127 of 1,213 MBCs (10.5%) encoded hallmark features of the IGHV3-30/14/Y motif, and a more restricted subset, defined by the IGHV3-30/14/[G/S-G-S/N-Y] motif, comprised 94 clones (7.8%) **(Figure S11)**. Within the IGHV3-30 lineage, these motif-bearing antibodies represented 38% and 28%, respectively. These findings indicate that antibodies targeting the Apex-B epitope are markedly expanded within the MBC compartment and may constitute a dominant class of S2-reactive, non-neutralizing antibodies, despite recognizing a single, highly focused epitope. In the study by Amanat *et al*., the vaccine study similarly demonstrated that, although vaccination markedly increased total antibody levels, neutralizing antibodies rapidly decreased in proportion compared to non-neutralizing antibodies^72^. Importantly, unbiased plasmablast sorting revealed that S2-specific antibodies accounted for approximately 20-52% of the total response across donors, highlighting that the S2 region naturally dominates the early response. Additionally, weak FP and SH responses in our study imply that S2-directed antibodies are largely focused on the S2-Apex. In addition, a previous study by Claireaux *et al*. reported that, even before the pandemic, healthy individuals possessed a surprisingly high frequency (0.086%) of SARS-CoV-2 spike-reactive naïve B cells, which was ∼30-fold higher than that observed for other unencountered antigens^17^. Among these antibodies, the IGHV1-69/IGKV3-11 pairing was strongly enriched and gave rise to non-neutralizing antibodies that bind the S2-Apex. The authors further observed a prominent representation of IGHV3-30 encoded antibodies, which likewise targeted the S2 subunit. Consistent with multiple studies, including our results, IGHV3-30 and IGHV1-69 are among the most strongly amplified IGHV genes after vaccination or infection. The high frequencies of both IGHV1-69/IGKV3-11 (as noted by Claireaux *et al*.) and IGHV3-30 subfamily Apex-B antibodies (as highlighted here) collectively reinforce the idea that the S2-Apex is a dominant node within the SARS-CoV-2 B-cell repertoire.

The S2-directed antibodies may arise either through recall of pre-existing seasonal CoVs induced memory B cells or through de novo activation of naïve B cells, as previously studied^17,22,73,74^. Notably, antibodies using IGHV1-69/IGKV3-11 or those targeting Apex-B exhibit germline-driven binding that does not require extensive SHMs, suggesting efficient recruitment from the naïve pool^17^. Given that Apex-B antibodies recognize *Sarbecovirus*-restricted sequence features, they are unlikely to be boosted by seasonal CoVs and thus most likely represent newly elicited responses. The persistence and expansion of these two major S2-Apex lineages into the memory B-cell compartment further indicate that this epitope is highly compatible with germinal-center selection, supporting its role as a recurrent and immunodominant target^17,71^.

Many S2-Apex binding antibodies, including those directed against Apex-B, have been reported as non-neutralizing^17–20,22^. For example, public clonotypes such as IGHV4-59/IGKV3-20 (M15) and IGHV1-69/IGKV3-11 engage structurally occluded regions but remain non-neutralizing, yet these antibody classes are markedly expanded following vaccination or natural infection^17,20^. These findings suggest that S2-Apex cryptic epitopes can act as immunological decoys, eliciting dominant antibody responses without providing protective benefit. Similar phenomena have been well described in the human immunodeficiency virus (HIV) field, where analogous strategies such as stabilizing the V3 loop, reducing CD4-binding affinity, or masking glycans were required to shift immunity away from decoy epitopes and toward protective determinants^75^.

It remains unclear how antibodies recognize the S2-Apex while the epitope is masked by the S1 domain. Although previous studies have suggested that conformational breathing motions can transiently expose this region^65,66^, direct structural evidence is still lacking. Even if breathing motions expose the Apex-B site sufficiently, how S1 remains associated with cleaved spikes after antibody binding remains an open question, as shown in Figures 6F and H. Our findings suggest that Apex-B antibodies preferentially engage spike conformations in which S1 is loosened or partially pre-shed, rather than actively inducing opening of the intact trimer. Notably, when the furin cleavage site is preserved, a substantial fraction of spikes occupies this destabilized state, even when the overall protein is stabilized. The restricted capacity of S2-Apex antibodies to engage the spike in its fully intact form may help explain their limited protective effect, despite their ability to elicit strong Fc-mediated functions such as antibody-dependent cellular cytotoxicity (ADCC)^17,18,22^. Studies by Changrob *et al*.^22^ and Johnson *et al*.^18^ reported the S2-Apex binding antibodies R125-61 and 54054-5, respectively. Both antibodies exhibited strong Fc-mediated effector functions. However, R125-61 showed only weak neutralizing activity and limited protective efficacy, whereas 54054-5 exhibited neither neutralizing activity nor measurable protection in vivo. Interestingly, Johnson *et al*. reported that abrogation of Fc effector function for 54054-5 resulted in a modest increase in protective activity. Rao *et al*.^20^ also described an antibody that binds near the S2-Apex region, engaging the spike trimer interface from the interior, which similarly lacked neutralizing and protective activity. In addition, although other S2-binding antibodies have been described, their protective efficacy has not been evaluated, and they have uniformly been reported as non-neutralizing^17,21^. To our knowledge, no S2-Apex binding antibody has been reported to confer meaningful antiviral protection. Therefore, although the protective efficacy of Apex-B binding antibodies was not directly evaluated in this study, prior evidence suggests that Fc-mediated effector functions of Apex-B binding antibodies are unlikely to translate into meaningful antiviral protection. Nevertheless, further studies will be required to directly assess whether Apex-B binding antibodies can contribute to protection in vivo.

A recent study reported on an antibody targeting the Apex-B site^19^. The antibody described, 6C10, shares the same epitope as COV2-2509 but utilizes IGHV1-2 and a 20-residue CDRH3. These characteristics differ from the IGHV3-30 antibodies observed in this study, and IGHV1-2 antibodies are not as frequently represented as those using IGHV3-30. Structural comparison of the two antibodies showed that, while 6C10 exhibits the same HC-dominant binding as COV2-2509, the orientations of the HC and LC are flipped relative to one another **(Figure S12A)**. Moreover, 6C10 covers a broader epitope than COV2-2509 **(Figure S12B)**. Thus, although the two antibodies share an overlapping epitope, they do not bind in an identical manner, and 6C10 is clearly distinct from the IGHV3-30 encoded Apex-B binding antibodies identified in this study. Additionally, that study showed that 6C10 did not confer significant protection in an in vivo mouse model, providing an example in which antibodies recognizing the Apex-B epitope do not translate into protective efficacy.

In conclusion, we demonstrate that the S2-Apex constitutes the most immunodominant region of the S2 subunit and provide the first characterization of Apex-B targeting IGHV3-30/14/Y-featured antibodies. These antibodies emerge as the most prominent clonotypes within the coronavirus antibody repertoire and exhibit convergent, public features. Although this epitope appears to preferentially elicit non-neutralizing antibodies, further studies are required to determine how selective depletion of S2-Apex-directed antibodies at the polyclonal level influences overall neutralization and protective activity. Together, these findings highlight the S2-Apex as a critical determinant of immunodominance and provide a framework for precision antigen design strategies aimed at eliciting more effective antiviral antibody responses in next-generation coronavirus vaccines.

## RESOURCE AVAILABILITY

### Lead contact

Further information and requests for resources and reagents should be directed to and will be fulfilled by the lead contact, Andrew B. Ward

### Materials availability

Stabilized Spike constructs are available upon request.

### Data and code availability

All data needed to evaluate the conclusions of this study are present in the paper and/or the Supplementary Information. Cryo-EMPEM data have been deposited in the Protein Data Bank (PDB) and the Electron Microscopy Data Bank (EMDB) and are publicly available as of the date of publication. PDB accession codes are 9Z80, 9ZT5, 9ZT6, 9ZT7, 9ZT8, and 10MU. EMDB accession codes are EMD-73884, EMD-75295, EMD-74737, EMD-74738, EMD-74739, EMD-74740, EMD-74741, EMD-74742, and EMD-74743. Ns-EMPEM data have been deposited in the EMDB under accession codes EMD-75193 and EMD-75194. Detailed information for each accession code is provided in Table S2. Any additional information required to reanalyze the data reported in this paper is available from the lead contact upon request. No code was generated in this study.

## ACKNOWLEDGEMENTS

We thank Hannah L. Turner, Anant Gharpure, and Charles A. Bowman for technical support with electron microscopy. We also thank Jeanne Matteson and Beverly Ellis for their contributions to mammalian cell culture. We are deeply appreciative of the generosity and long-term engagement of our study participants, whose continued involvement made this research possible. Special thanks go to the Personalized Virology Initiative team for sharing biospecimen and metadata from study participants as well as to Carter Williams, Morgan van Kesteren and Hyun Min Kang for excellent technical assistance. The authors would like to thank Drs. Nicole Doria-Rose, Lingshu Wang and Qiong Zhou, Humoral Immunology Core Vaccine Research Center, NIH for the VRC7568_SARS-CoV-2 S_D614G (VRC_LW, Lot U794QFH180-2, pHR’ CMV Luciferase (VRC-5601) plasmids, ; Dr. Frank Kirchhoff, University of Ulm, for the pCG_SARS-CoV-1 S (humanized) C-V5 IRES GFP plasmid, and Dr. Paul Bieniasz, Rockefeller University, for pNL Gag Pol plasmid.

This research was supported in part by the National Institute of Allergy and Infectious Diseases of the National Institutes of Health under Award Numbers P01 AI172531 (to A.B.W., F.K., and V.S.) and R01 AI190286 (to A.B.W. and I.A.W.), and by institutional funds provided to the Mount Sinai Center for Vaccine Research and Pandemic Preparedness (to F.K. and V.S.).

## AUTHOR CONTRIBUTIONS

Conceptualization, S.P. and A.B.W.; Supervision, A.B.W., I.A.W., S.B., and M.Y.; Cryo-EMPEM and ns-EMPEM data collection and analysis, S.P.; BCR sequence analysis, S.P.; Study participant recruitment and collection of human biospecimen, K.S., C.G., and V.S.; Virus and cell-based assays, J.M., N.O., LCFM and A.A.; Protein production and plasma processing, S.P., J.C., and K.S.; Material resources, A.B.W., F.K., V.S., and I.A.W.; Writing-original draft and editing, S.P., A.B.W., and S.B. All authors reviewed and approved the final version of the manuscript.

## METHODS

### Human subjects

The plasma used were sourced from an IRB approved observational research study protocol [STUDY-16-01215/IRB-16-00971] that collects biospecimen from adults with confirmed or potential exposure to viruses in New York City. The study was reviewed and approved by the Program for the Protection of Human Subjects at the Icahn School of Medicine at Mount Sinai. All study participants provided written informed consent prior to data and sample collection including biospecimen banking/sharing.

Plasma from nine adult study participants (7 women, 2 men) were selected based on SARS-CoV-2 immune histories (e.g., number and sequence of vaccination and infections). Two participants had not received a COVID-19 vaccine prior to the collection of the plasma used in this study while the other seven had completed their initial COVID-19 immunizations (Pfizer BNT162b2: 4, Moderna mRNA-1273: 3). Plasma from five participants was collected after they received one (N: 5) or two (N: 2) booster immunizations. In addition, three of nine participants had no documented SARS-CoV-2 infection, while six had one or two documented, symptomatic but mild COVID-19 manifestations (no medical interventions such as hospitalization required).

### Expression and purification of spike proteins

For stabilized S2 proteins (S2-E69, S2-HP, S2-HP-2W, S2_W4_, and S2_W4.1_; Table S1), genes encoding the S2 proteins were cloned into a pcDNA3.4 vector containing either a C-terminal His-Tag or a Twin-strep tag. The expression plasmids were transfected into Expi293F cells (Thermo Fisher Scientific). After 6 days, cells were harvested, and supernatants containing the proteins were clarified by centrifugation and incubated with Ni-nitrilotriacetic acid (NTA) resin (Cytiva) or Strep-Tactin XT resin (IBA Lifesciences) for 1h at 4℃. For Ni Sepharose excel resin (Cytiva), the resin was washed with a solution of 40 mM imidazole, 20 mM Tris, pH 7.5, and 150 mM NaCl. After the wash step, proteins were eluted using 300 mM imidazole, 20 mM Tris, pH 7.5, and 150 mM NaCl. For Strep-Tactin XT resin, the resin was washed with a solution of 20 mM Tris, pH 7.5, and 150 mM NaCl, followed by proteins were eluted with a buffer 1xBXT (IBA Lifesciences). The S2 proteins were finally purified by size-exclusion chromatography using a HiLoad 16/600 Superdex 200 (Cytiva) column.

For full-length spike proteins, we generated the ancestral SARS-CoV-2 spike with six stabilizing prolines (CoV2-HP-S, Wuhan-Hu-1 strain) and several modified constructs: one containing the native S1/S2 furin cleavage site (residues 682-685; S-HP-RRAR), one with a GSAS substitution at the cleavage site (S-HP-GSAS), and two containing additional inter-protomer disulfide bonds (S-HP-Mut7-GSAS and S-HP-Mut7-RRAR) as described previously^50,64,76^.

Variant constructs including Omicron (B.1.1.529), BA.4/5, BA.1.5, JN.1, and KP.3.1.1 were generated by introducing the respective mutations into the S-HP background. The OC43 spike construct comprised residues 1-1287 (GenBank AIL49484.1) with stabilizing proline substitutions at positions 1079 and 1080. The SARS-CoV spike (CoV-2P-S) construct was generated from residues 1-1196 of the Tor2 strain (GenBank AAP41037.1) with stabilizing proline substitutions at positions 968 and 969.^64^ The HKU1 spike includes residues 1 to 1276 from isolate N5 (GenBank Q0ZME7). Briefly, the OC43, HKU1, and CoV-S spike proteins were purified using the N-terminal His-Tag with cOmplete resin (Milipore Sigma) and eluted with 250 mM imidazole. The eluates were buffer-exchanged into Tris-NaCl buffer (25 mM Tris, 500 mM NaCl, pH 7.4) and subsequently purified on a Superose 6 Increase 10/300 column (Cytiva). For CoV2-S spike proteins, purification was carried out using Strep-Tactin XT high-capacity columns (IBA Lifesciences). Proteins were eluted with BXT buffer (IBA Lifesciences) and further purified using a Superose 6 Increase column.

For monoclonal Fab and IgG purification (mFab and mIgG), all mFabs and mIgGs were expressed using the Expi293 cell system. The plasmids for heavy and light chains of each antibody were co-transfected, and supernatants were harvested after 6 days. The antibodies were purified using CaptureSelect XL CH1 resin (Thermo Fisher Scientific), followed by further purification by size-exclusion chromatography (SEC) in 20 mM Tris (pH 7.5), 150 mM NaCl buffer using a HiLoad 16/600 Superdex 200 (Cytiva) column.

### Purification of human plasma IgG and Fab

For IgG isolation, 5 ml of human plasma was incubated with 5 ml of pre-washed protein G resin (Cytiva) at 4°C for 2 days with gentle rotation. After incubation, the resin was washed three times with PBS and subsequently eluted using 2.0 M glycine buffer (pH 2.5). The eluate was immediately neutralized with 1 ml of 1 M Tris-HCl (pH 9.0) and buffer-exchanged into PBS using a 100-kDa cutoff Amicon ultrafiltration device. For IgG preparation, undigested IgG was further purified using a Superdex 200 16/600 column (Cytiva). For Fab preparation, purified polyclonal IgG was digested with papain (Thermo Fisher Scientific) at 37°C for 5 hours, using 40 µg of papain per 1 mg of IgG, in a digestion buffer composed of 100 mM Tris-HCl (pH 8.0), 2 mM ethylenediaminetetraacetic acid (EDTA), and 10 mM L-cysteine. Reaction was quenched using 0.03 M iodoacetamide. Following digestion, the reaction mixture was separated from the papain resin and buffer-exchanged into phosphate-buffered saline (PBS). Undigested IgGs were removed by size-exclusion chromatography on a Superdex 200 16/900 column (Cytiva).

### Monoclonal antibody expression

In this study, all monoclonal antibodies were cloned and expressed using the TurboCHO antibody expression service (GenScript). For cloning, variable heavy (VH) sequences were cloned into human IgG1 constant heavy chain expression vectors, while variable light (VL) sequences were cloned into either human kappa or lambda light chain expression vectors.

### Differential scanning calorimetry (DSC)

Samples were prepared at a concentration of 0.4 mg/mL in Tris-buffered saline (TBS) buffer (pH 8.0). Prior to each measurement, both sample and reference cells were thoroughly washed twice with TBS buffer and allowed to equilibrate. Approximately 300 μL of sample solution was loaded into the sample cell, while the reference cell was filled with an equal volume of TBS buffer. DSC measurements were performed using a temperature program that started at 20°C and was followed by heating to 100°C at a scan rate of 90°C/h. Data processing included buffer subtraction using the reference cell baseline, normalization, and baseline correction. All data analysis was performed using Origin 7.0 software. Melting temperatures (Tm) were determined from the peak maximum of the heat capacity curves.

### Ns-EMPEM sample data collection and processing

Fab-S2_W4.1_ complexes were generated by incubating 15 μg of S2 protein with 1 mg of purified polyclonal Fabs overnight at 4°C with gentle rotation. To remove excess unbound Fabs, the samples were washed five times with TBS buffer using a 500 μL 100-kDa cutoff Amicon ultrafiltration device. The final concentrated complex was adjusted to approximately 0.01 mg/mL and immediately used for preparing EM grids. For making grids, 3 μL of complex sample was applied to carbon-coated 400-mesh copper grids (made in-house) and stained with 2% (w/v) uranyl formate for an initial 30 seconds, followed by an additional 60 seconds.

Grids were imaged at 200 kV on a TFS TALOS F200C system (Thermo Fisher Scientific) using a FEI Ceta (4k × 4k) camera. Micrographs were collected using the EPU program, and post-processing was conducted using Relion 4.0^77^. Particles were picked using auto-picking and subjected to 2D classification. The complex particle stacks from 2D classification were selected and subjected to iterative 2D classification (2-3 rounds), followed by 3D classification to sort classes based on different Fab specificities. ChimeraX was used to visualize the composite map, which included segmented unique Fab classes and S2 protein maps^78^

To calculate the EMPEM magnitude for each Apex-A and Apex-B epitope, Fab-bound 2D classes were selected and subjected to initial 3D refinement using Relion 4.0 with C1 symmetry for Apex-A and C3 symmetry for Apex-B. Particles corresponding to Apex-B were C3 symmetry-expanded, whereas Apex-A particles were not expanded because the epitope is located on the symmetry axis. Epitope-bound particles were then classified by focused 3D classification using a 40 Å-diameter spherical mask centered on the epitope site as described previously^79^. The EMPEM magnitude for each epitope was calculated by dividing the number of final epitope-bound particles by the total number of particles used in the initial 3D refinement.

### Cryo-EMPEM sample preparation

For cryo-EMPEM analysis, S2_W4.1_ protein of 50 μg was complexed with 5 mg of purified polyclonal Fabs from each donor and incubated overnight at 4°C with gentle rotation. The resulting complexes were purified using a Superose 6 Increase 10/300 column (Cytiva) equilibrated in buffer containing 20 mM Tris (pH 7.5) and 150 mM NaCl. The purified complex at a concentration of 0.5-0.7 mg/ml was mixed with 0.5 μl of octyl-β-glucoside (OβG) detergent to achieve a final concentration of 0.1% (w/v). The sample was then immediately applied to glow-discharged 1.2/1.3 300-mesh UltraAuFoil grids and plunge-frozen using a Vitrobot Mark IV (Thermo Fisher Scientific).

### Cryo-EMPEM data collection and image processing

All cryo-EMPEM datasets were collected on a 200 kV Glacios (Thermo Fisher Scientific) equipped with a Falcon 4i direct electron detector. Automated data collection was carried out using EPU software (Thermo Fisher Scientific) at a nominal magnification of 190,000× with a pixel size of 0.718 Å. Movies were recorded with an exposure dose of approximately 45 e⁻/Å² and a defocus range of −0.8 to −1.8 μm. All dataset movies were motion-corrected with dose weighting and CTF estimation was performed using CryoSPARC Live.

All dataset processing was conducted using CryoSPARC^80^. Particles were picked using blob picker or template picker. Clean particle stacks were selected through multiple rounds of 2D classification and used to generate reference volumes via ab initio reconstruction, followed by heterogeneous refinement. After heterogeneous refinement, different antibody-bound conformations were sorted using focused classification protocols described previously^38^. The epresentative data processing workflow is summarized in Figure S13, and detailed data collection parameters are provided in Table S2.

### Cryo-EM sample preparation for S2_W4.1_ apo and S2_W4.1_-COV2-2509 Fab complex

For cryo-EM analysis, the S2_W4.1_ apo protein at 3 mg/mL was mixed with 0.5 μl of CHAPSO (3-[(3-cholamidopropyl)dimethylammonio]-2-hydroxy-1-propanesulfonate, final concentration 4 mM), while the S2_W4.1_-COV2-2509 Fab complex was prepared at 0.5 mg/ml. Samples (3 μl each) were applied to glow-discharged 1.2/1.3 300-mesh UltraAuFoil and vitrified by plunge-freezing in liquid ethane using a Vitrobot Mark IV (Thermo Fisher Scientific) maintained at 20°C and 100% humidity with a blot time of 3 seconds.

### Cryo-EM data collection and image processing for S2_W4.1_ apo and S2_W4.1_-COV2-2509 Fab complex

All cryo-EM datasets were collected on a 200 kV Glacios (Thermo Fisher Scientific) equipped with a Falcon 4i direct electron detector. Data collection steps followed the same protocols as described above for the cryo-EMPEM datasets. For S2_W4.1_ apo image processing, particles underwent multiple rounds of 2D classification to generate clean particle stacks, followed by ab initio reconstruction to generate initial reference models. Heterogeneous refinement was performed to separate different conformational states. The final model was obtained through global CTF refinement and reference-based motion correction, followed by non-uniform refinement with C3 symmetry imposed. For the S2_W4.1_-COV2-2509 Fab complex, initial processing followed the same workflow as the apo structure. Subsequently, focused 3D classification was applied to the Fab region to isolate Fab-bound particles. The selected particles were then subjected to C3-symmetry expansion and local refinement with C1 symmetry to obtain the final structure. Data collection parameters are provided in Table S2.

### Model building and refinement

For cryo-EMPEM datasets, the S2_W4.1_ structure was initially built using the S2 subunit of PDB ID 6XR8 as a template. Fab coordinates were modeled as poly-alanine backbones based on the COV2-2509 Fab, which was initially generated with ABodyBuilder2^81^. The first round of refinement was performed using Rosetta relaxed refinement^82^, followed by iterative cycles of manual model building in Coot and real-space refinement in Phenix^83,84^. EMRinger and MolProbity scores were calculated after each Rosetta refinement round to guide model quality assessment^85,86^. For Cryo-EM datasets, the S2_W4.1_ structure obtained from cryo-EMPEM was used as the starting model, while the COV2-2509 Fab was initially modeled with ABodyBuilder2. Subsequent refinements were carried out following the same iterative procedure described above. All structural figures were generated using ChimeraX^78^. Final model maps, validations, and statistics are summarized in Table S2 and Figures S5.

### Sequence-to-structure analysis (STS) with ModelAngelo (MA)

The cryo-EMPEM maps corresponding to the Apex-B epitope antibody class from donors 1, 2, 3, and 8-T2 were used for STS analysis. This analysis basically followed our previous report^42^. Briefly, predicted sequences for the Fab-related map density in each analysis were generated using the “model_angelo build_no_seq” command from MA. The multiple fragmented Fab sequences obtained from MA were structurally aligned and compared with known antibody structures using the accompanying output.cif file to determine their proper sequence order. Based on this determined order, the fragmented .hmm profiles were then assembled into a single integrated heavy chain .hmm profile **(Figure S6A)**. The integrated .hmm file was then compared against the CoV-AbDab database containing 12,918 antibodies using MA’s hmm_search function, and antibodies with the lowest E-value scores were selected.

### ELISA analysis

To assess epitope-specific antibody titers in longitudinal samples from donors 8 and 9, streptavidin-coated 96-well plates (Thermo Scientific) were coated with twin-strep-tagged proteins (S2_W4.1_N*FP, S2_W4.1_ΔSH) starting at 0.2 mM and biotin-Ahx-modified (SH_pep_, FP_pep_; Genscript) starting at 3 mM were prepared in 3-fold serial dilutions in buffer containing 20 mM Tris pH 7.5, 150 mM NaCl, 0.1% BSA, and 0.05% Tween-20 (TBST buffer), and coated on streptavidin-coated 96-well plates at 100 μl per well for 2 hours at room temperature (RT). After three washes, plasma samples from each donor were diluted 1:400 in TBST buffer and incubated in 100 μl per well for 2 hours at RT. After incubation, plates were washed three times with TBST buffer and incubated with anti-human IgG-Horseradish Peroxidase (HRP) secondary antibody (Invitrogen) for 1 hour. Following three additional washes with TBST buffer, 3,3’,5,5’-Tetramethylbenzidine (TMB) substrate was added for color development, and absorbance was measured at 450 nm using a spectrophotometer.

For epitope-specific antibody titer comparison across multiple donors with different immune histories (Infection [I], Vaccination [V], and Infection+Vaccination [VI]), streptavidin-coated 96-well plates were coated with 0.2 mM of each protein antigen (S2_W4.1_ΔSH, Full-spike: CoV2-S-RRAR) and 3 mM peptide antigen (SH_pep_, FP_pep_) using the same coating procedure described above. Plasma samples from each donor were diluted 1:1000 in TBST buffer and incubated as a single-point well for 2 hours at room temperature. Following the same washing and detection procedures, OD_450_ signals were measured for each antibody response. All graphical analyses and statistical comparisons were performed using GraphPad Prism 10.

### CoV-AbDab and BCR sequencing analysis

CoV-AbDab sequence analysis was performed using 11,180 human B cell-derived sequences selected from the total database of 12,918 sequences. Non-human sequences and those with chimeric, computational, engineered, humanized, mouse, phage display, yeast display, or unknown origins were excluded. Selected sequences were categorized into three epitope groups based on their binding annotations: S1 domain (S; RBD, S; NTD, S; S1, N, S; NTD-SD1, S; Possibly RBD, N; NTD, S; S1 non-RBD), S2 domain (S; S2 Fusion Peptide, S; S2, S; S2 (HR2 Peptide), S; non-S1, S; S2 Fusion Peptide, S; S2’ Cleavage Site/Fusion Peptide, NTD, S; S2 Stem Helix), and unknown epitopes (S; Unk, S; non-RBD Unknown). BCR sequencing analysis was performed using previously reported data from Kim *et al*.^53^. Only heavy chain sequences were analyzed, with analysis restricted to S-positive sequences. Both bulk sequencing and 10x Genomics single-cell sequencing data were utilized for the analysis. In the analysis, the IGHV3-30 family included IGHV3-30, IGHV3-30-3, and IGHV3-30-5. All statistical analyses and data visualization were performed using GraphPad Prism 10 and R. Alluvial diagrams were created using the ggalluvial package^87^, and sequence logo plots were generated using WebLogo3^88^.

### Frequency analysis of IGHV3-30/14/Y-feature antibodies

For frequency analysis of IGHV3-30/14/Y-feature antibodies across donor groups with different exposure histories (healthy donors, SARS-CoV-2 infection, and vaccination), BCR sequencing data were obtained from the OAS database^60^ and the dataset reported by Briney *et al*.^59^. The healthy donor group included 10 subjects from the Briney *et al*. dataset (https://github.com/briney/grp) and 11 subjects from unpaired BCR sequences in the OAS database. The infection group consisted of 70 subjects derived from unpaired sequences in the OAS database. The vaccination group comprised 50 subjects from the paired sequences dataset in the OAS database. For each subject dataset, unique BCR clones were identified by defining clonotypes as sequences with an exact match of V gene, J gene, and CDR3 amino acid sequence. Clonotypes carrying the IGHV3-30/14/[G/S-G-S/N-Y] motif were enumerated, and their frequency was calculated relative to the total number of unique BCR clonotypes within each dataset. Frequencies were normalized and reported as counts per million unique BCR clonotypes.

### Size-exclusion chromatography analysis

To determine the binding property between COV2-2509 and the full-length spike protein, 200 µg of S-HP-GSAS or S-HP-RRAR trimer was incubated with COV2-2509 IgG at a 1:1 molar ratio (spike monomer:IgG) for 2 h at 4°C. Following incubation, the mixtures were subjected to size-exclusion chromatography (SEC) on a Superose 6 Increase 10/300 GL column (Cytiva) pre-equilibrated with 20 mM Tris (pH 7.5) and 150 mM NaCl. Chromatograms were recorded and analyzed using UNICORN 6.3 software. Fractions corresponding to the eluted complexes were collected and evaluated by SDS-PAGE to confirm complex formation.

### Biolayer interferometry (BLI) analysis

For assays measuring the interaction between S2_W4.1_ and antibodies, Ni-NTA biosensors (Sartorius) were pre-hydrated in kinetic buffer composed of 1× PBS (pH 7.4) supplemented with 0.1% bovine serum albumin (BSA) and 0.05% Tween-20, followed by equilibration to establish a stable baseline. His-tagged S2_W4.1_ protein (10 µg/mL) was immobilized onto the biosensor surface, and sensors were washed in kinetic buffer to remove unbound protein. Association was measured by exposing the sensors to antibody solutions (25 or 50 µg/mL) for 300 s, followed by dissociation in kinetic buffer for an additional 300 s. For assays measuring the interaction between COV2-2509 and full-length spike proteins, anti-human IgG Fc Capture (AHC) biosensors (Sartorius) were used to immobilize COV2-2509 IgG (10 µg/mL). The subsequent steps were identical to those described above, with spike proteins (50 µg/mL) allowed to associate for 300 s and dissociate for 300 s in kinetic buffer. For competitive binding analysis between COV2-2509 and other antibodies targeting the Apex-B epitope, His-tagged S2_W4.1_ protein was immobilized onto Ni-NTA biosensors as described above. After washing to remove unbound protein, sensors were first incubated in wells containing COV2-2509 antibody (50 µg/mL) for 300 s to achieve saturation binding. The sensors were then transferred to wells containing the second antibody and incubated for an additional 300 s to measure association in the presence of pre-bound COV2-2509. Data were reference-subtracted and analyzed with ForteBio Data Analysis software.

### Cell culture

Vero.E6 cells expressing transmembrane protease serine 2 (TMPRSS2) (BPS Biosciences, cat. no. 78081) were maintained in Dulbecco’s modified Eagle’s medium (DMEM; Gibco, cat. no. 11965092) supplemented with 10% heat-inactivated fetal bovine serum (FBS; Gibco, cat. no. A5256801), 1% minimum essential medium with non-essential amino acids (Gibco, cat. no. 11140050), 100 U/mL penicillin and 100 μg/mL streptomycin (Gibco, cat. no. 15140122), 100 μg/mL normocin (InvivoGen, cat. no. ant-nr), and 3 μg/mL puromycin (InvivoGen, cat. no. antpr). HEK-293T cells (ATCC, CRL-3216) and HEK 293T-ACE2-TMPRSS2 (BEI Resources # NR-55293) were maintained in Dulbecco’s modified Eagle medium (Corning, 10-013-CV) supplemented with 10% heat-inactivated FBS (GeminiBio, 100-106), 10% L-Glutamine (Corning, 25-005-CI), and 10% penicillin-streptomycin (Corning, 30-002-CI) and cultured in a 5% carbon dioxide atmosphere at 37°C.

### Multicycle SARS-CoV-2 neutralization assays

Neutralizing antibody titers against SARS-CoV-2 (USA-WA1/2020; BEI Resources, NR-52281) were determined using a multicycle microneutralization assay performed in a BSL-3 facility. Vero.E6-TMPRSS2 cells (2 × 10⁴ cells/well) were seeded in 96-well plates 24 h prior to infection. S2 monoclonal antibodies were serially diluted 3-fold starting at 30 µg/mL in MEM (1×) and incubated with 10,000 TCID₅₀/mL of virus for 1 h at room temperature. Remdesivir (Medkoo Bioscience Inc.) was used as an internal control and was serially diluted alongside the monoclonal antibodies. Virus-antibody mixtures were transferred to the cell monolayers and incubated for 1 h at 37°C. After adsorption, the inoculum was removed and cells were overlaid with MEM (1×) containing 2% FBS, followed by incubation for 48 h at 37°C. Cells were fixed with 10% formaldehyde overnight at 4°C, permeabilized with 0.1% Triton X-100, and blocked in PBS containing 3% BSA. Biotinylated anti-SARS-CoV nucleoprotein mAb 1C7C7^89^ (1 µg/mL) in PBS plus 1 % BSA was applied for 2 h at RT. Cells were washed with 200 μl/well of 1 x PBS twice and 100 μl/well of HRP-conjugated streptavidin (Thermo Fisher Scientific) diluted in PBS plus 1 % BSA were added at a 1:2000 dilution for 1 h at RT. Cells were washed twice with PBS, and 100 μl/well of Sigmafast OPD (Sigma-Aldrich) were added for 10 min at RT, followed by addition of 50 μl/well of a 3 M HCl solution (Thermo Fisher Scientific). Absorbance was measured at 490 nm. Fifty-percent inhibitory dilution (ID₅₀) values were calculated by non-linear regression with the upper and lower plateaus constrained to 100% and 0%, respectively. Two independent experiments were performed.

### Pseudovirus production and single cycle entry assays

Coronavirus spike-pseudotyped lentiviral particles carrying a luciferase reporter were generated by transient transfection of HEK293T cells as previously reported^90^. Briefly, 7 × 10⁶ HEK 293T cells were transfected with plasmids encoding spike (SARS-CoV-2 S_D614G or SARS-CoV-1 S), lentiviral Gag-Pol, and luciferase reporter were co-transfected at a 1:16:16 ratio using polyethylenimine (PEI). Filtered supernatants aliquoted, and stored at −80°C.

Pseudovirus-based SARS-CoV-1/2 neutralization assays were performed using 293T-ACE2-TMPRSS2 cells as previously reported^90^. S2 Apex-B antibodies were serially diluted (initial 1:10 dilution followed by six 3-fold steps) in culture medium containing SARS-CsoV-1 or SARS-CoV-2 spike-pseudotyped luciferase lentiviral particles and incubated for 45 min at 37 °C, 5% CO₂. The antibody/pseudovirus mixtures were added to the target cells, followed by incubation at 37 °C, 5% CO₂ for 48 hours. Luciferase activity was measured using the Luciferase Assay System (Promega, E4530) on a VICTOR3™ multilabel plate reader (PerkinElmer). Neutralization titers (ID₅₀) were calculated by determining the antibody concentration yielding >50% inhibition using a variable-slope logistic model fitted to normalized luminescence data in Prism 10.

### ADCC reporter assay

Vero.E6 cells were infected with NDV-HexaPro-S^91^ at a multiplicity of infection (MOI) of 2 and incubated for 72 hours. Following incubation, the media was removed and 25 μL of RMPI-1640 media supplemented with 4% low-IgG FBS was added. 25 μL of serially diluted monoclonal antibodies (30-0.06 μg/mL) were added to the plate in duplicates. ADCC Bioassay Effector Cells (Promega, G7102) were added (1.5x10^5^ cells/well) and the plates were incubated for 6 hours. 75ul of Bio-Glo luciferase reagent from the Bio-Glo™ Luciferase Assay System (Promega, G7940) were added to each well and the luminescence was read using a Synergy 4 (BioTek) plate reader. The area under the curve (AUC) was calculated using GraphPad Prism 10. The COV44-79 and COV89-22 antibodies served as S2-binding antibody controls.

