## Supplemental Figures and Tables for "The buried S2 apex of SARS-CoV-2 spike elicits an immunodominant germline-restricted public antibody response"

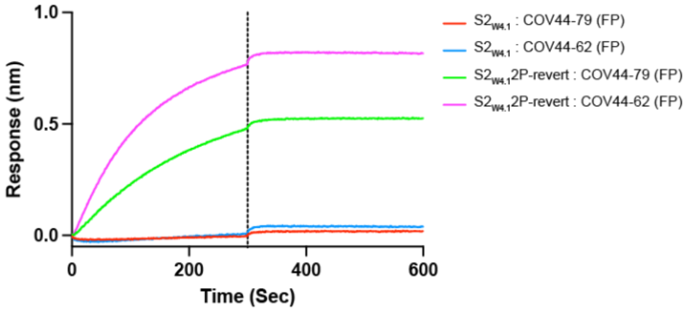


**Figure S1. Binding kinetics of FP-directed antibodies to S2 constructs measured by biolayer interferometry (BLI).** Reversion of the K986P and V987P (2P) mutations in $S2_{W4.1}$ ($S2_{W4.1}$2P-revert) enhanced binding to fusion peptide (FP) antibodies COV44-79 and COV44-62 compared to $S2_{W4.1}$.

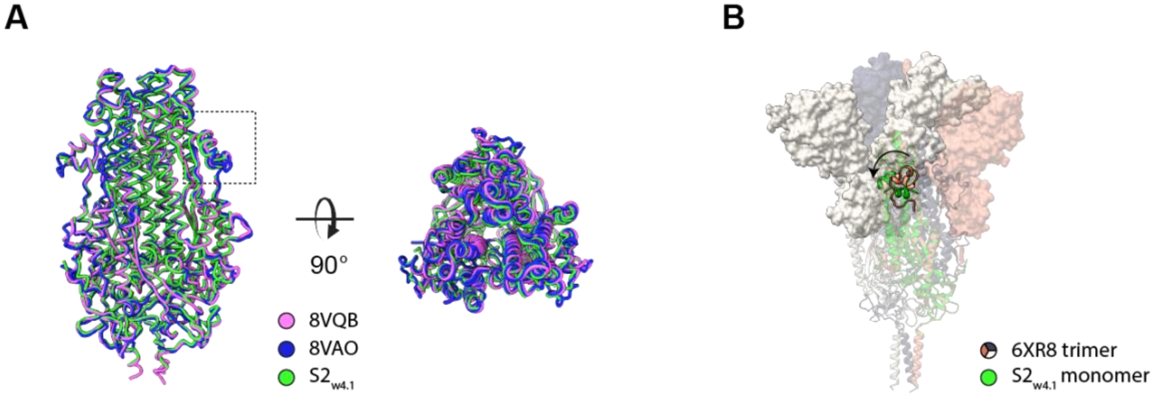


**Figure S2. Structural comparison of S2$_{W4.1}$ with previous SARS-CoV-2 S2 structures and full-length spike structure.** (A) Superposition of S2$_{W4.1}$ structure (green) with previously reported S2 subunit structures from PDB ID 8VQB (magenta) and 8VAO (blue). The dotted box highlights the region showing structural variations depicted in (B). (B) Structural comparison between S2$_{W4.1}$ and full-length spike structures. The only monomer of S2$_{W4.1}$ trimer is superimposed onto the full-length spike structure (PDB ID 6XR8). The S1 subunit of the full-length spike is depicted as a surface model, and the S2 subunits are represented in cartoon models. Residues 828-856, which show structural differences between the two structures, are highlighted while other regions are shown with reduced opacity. The arrow indicates the conformational transition of residues 828-856.

A

| | Total | Apex-A (magnitude) | Apex-B (magnitude) |
|---|---|---|---|
| donor1 | 13,189 | 1,084 (0.08) | 31,138 (2.36) |
| donor2 | 25,486 | 2,623 (0.10) | 95,928 (2.06) |
| donor3 | 46,625 | 17,301 (0.37) | 62,461 (2.45) |
| donor4 | 34,221 | 2,766 (0.08) | 83,922 (2.45) |
| donor5 | 77,573 | 32,736 (0.42) | 102,683 (1.32) |
| donor6 | 36,657 | 12,051 (0.33) | 70,912 (1.93) |
| donor7 | 11,331 | 5,780 (0.51) | 18,514 (1.63) |
| donor8-T1 | 5,501 | 722 (0.13) | 14,909 (2.71) |
| donor8-T2 | 263,773 | 110,504 (0.42) | 664,557 (2.52) |
| donor8-T3 | 112,306 | 11,232 (0.10) | 253,162 (2.25) |
| donor9-T1 | 44,713 | 33,314 (0.75) | 66,745 (1.49) |
| donor9-T2 | 17,440 | 16,867 (0.97) | 33,150 (1.90) |

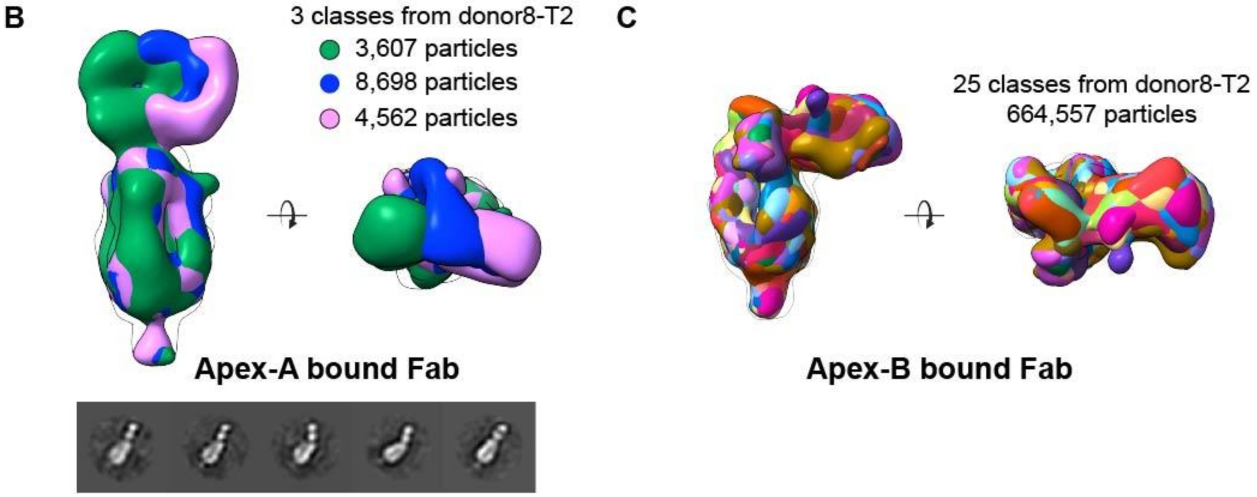


**Figure S3. Distinct Fab density patterns at the Apex-A and Apex-B epitopes revealed by ns-EMPEM.** (A) Per-donor particle counts assigned to the Apex-A and Apex-B epitopes used to calculate EMPEM magnitude. Particle numbers for Apex-B can exceed the total particles due to C3 symmetry expansion (see Methods) (B) Epitope-specific 3D classification of Apex-A bound Fab densities from ns-EMPEM analysis of plasma from Donor 9-T2. Three major 3D classes were identified, indicating heterogeneous Fab density and binding orientations at the Apex-A epitope. Representative 2D classes are shown below. (C) Epitope-specific 3D classification of Apex-B bound Fab densities from Donor 8-T2. Twenty-five 3D classes were identified from 664,557 particles, showing more uniform Fab density and higher occupancy at the Apex-B epitope.

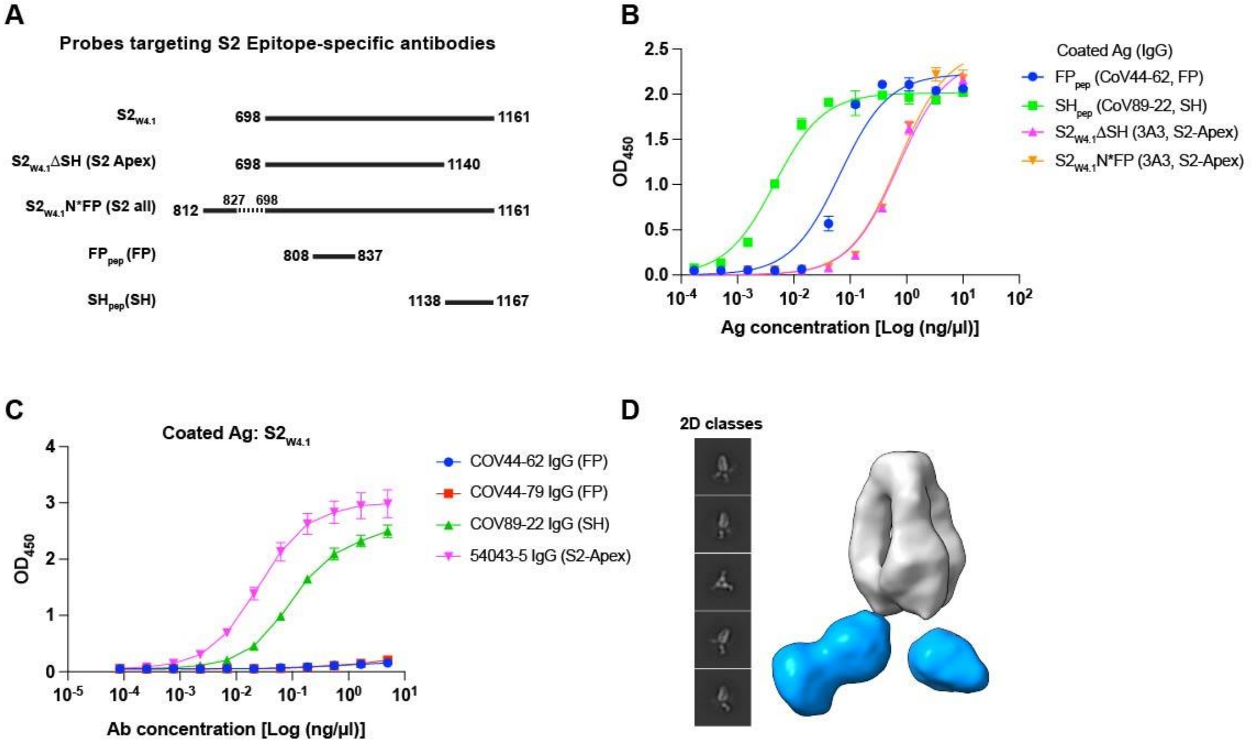


**Figure S4. SARS-CoV-2 S2 epitope-specific probe design and binding validation.** (A) Schematic representation of protein and peptide constructs designed as probes for targeting S2 epitope-specific antibodies. Numbers indicate residue positions in the SARS-CoV-2 spike sequence. (B) ELISA binding curves of epitope-specific monoclonal antibodies against the indicated probes. Antibody concentration was fixed at 10 µg/ml, while coated antigens were serially diluted. Data show similar $B_{max}$ saturation binding profiles, indicating that the relative antibody levels are comparable across different probes. Error bars indicate mean ± SD of technical replicates. (C) ELISA binding curves of fusion peptide (FP), stem helix (SH), and S2-Apex antibodies to the $S2_{W4.1}$ probe. S2-Apex (54043-5) and SH (COV89-22) antibodies showed strong binding, whereas FP-targeting antibodies (COV44-62 and COV44-79) exhibited negligible binding, confirming the lack of FP interaction in $S2_{W4.1}$. (D) Representative 2D class averages and ns-EM reconstruction of $S2_{W4.1}$ bound to the SH antibody (COV89-22). The Fab density is shown in blue, and the S2 trimer density in gray.

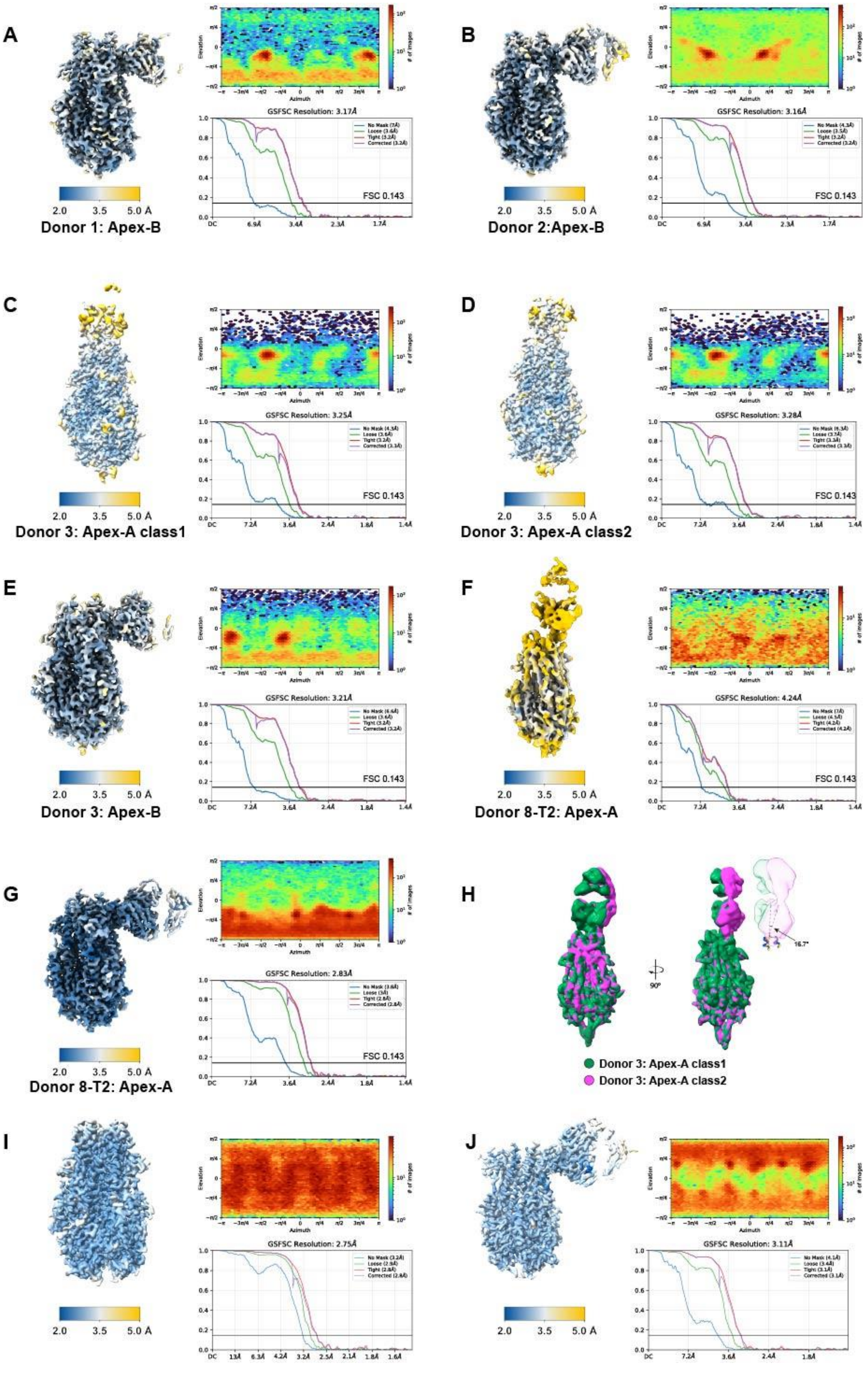

A
2.0 3.5 5.0 Å
Donor 1: Apex-B
GSFSC Resolution: 3.17Å
FSC 0.143
B
2.0 3.5 5.0 Å
Donor 2:Apex-B
GSFSC Resolution: 3.16Å
FSC 0.143
C
2.0 3.5 5.0 Å
Donor 3: Apex-A class1
GSFSC Resolution: 3.25Å
FSC 0.143
D
2.0 3.5 5.0 Å
Donor 3: Apex-A class2
GSFSC Resolution: 3.28Å
FSC 0.143
E
2.0 3.5 5.0 Å
Donor 3: Apex-B
GSFSC Resolution: 3.21Å
FSC 0.143
F
2.0 3.5 5.0 Å
Donor 8-T2: Apex-A
GSFSC Resolution: 4.24Å
FSC 0.143
G
2.0 3.5 5.0 Å
Donor 8-T2: Apex-A
GSFSC Resolution: 2.83Å
FSC 0.143
H
90°
16.7°
Donor 3: Apex-A class1
Donor 3: Apex-A class2
I
2.0 3.5 5.0 Å
GSFSC Resolution: 2.75Å
J
2.0 3.5 5.0 Å
GSFSC Resolution: 3.11Å

**Figure S5. Cryo-EMPEM data validation and Apex-A antibody class analysis.** (A-G) Local resolution maps (left), angular distribution (right top), and FCS curve (right bottom) analyses are shown for each reconstruction. Resolution map colored based on the resolution estimated at FSC 0.143, gold-standard FSC (GSFSC) (H) Comparison of Apex-A bound Fab classes from Donor 3. Two distinct classes are observed, with a relative angular difference of ~16.7°, displaying structural heterogeneity in antibody engagement at the Apex-A epitope. (I and J) Local resolution maps, angular distributions, and FCS curves for $S2_{W4.1}$ apo and $S2_{W4.1}$-COV2-2509 Fab complex, respectively.

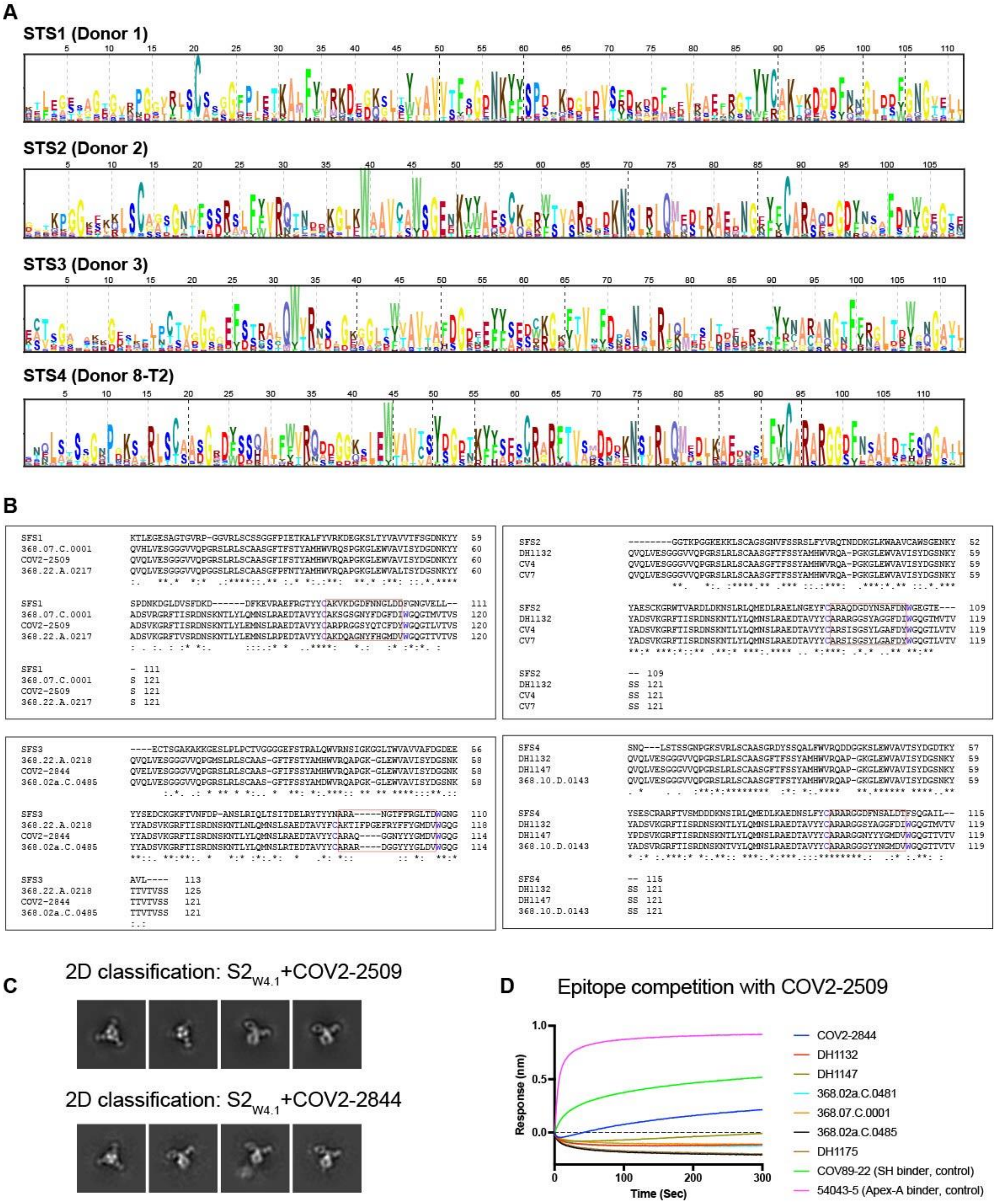


**Figure S6. Sequence-to-structure (STS) analysis and epitope validation.** (A) Sequence logo plots derived from hidden Markov models (HMMs) of each STS dataset, ordered according to the reference heavy-chain sequence. Logos were generated using Skylign[92]. (B) Sequence alignment of representative HMM-derived sequences with their closest matching antibodies from CoV-AbDab. CDRH3 positions are highlighted with red boxes, and the flanking residues Cys92 and Trp103 are marked in blue. (C) Representative 2D classes from ns-EM validating S2 binding of COV2-2509 and COV2-2844 to the same epitope. (D) Competitive biolayer interferometry (BLI) assay. Sensor chips were pre-incubated with COV2-2509 before antibody association. All tested antibodies showed complete competition with COV2-2509, except COV2-2844, which exhibited partial competition. Ns-EM analysis further confirmed that COV2-2844 binds to the Apex-B epitope.

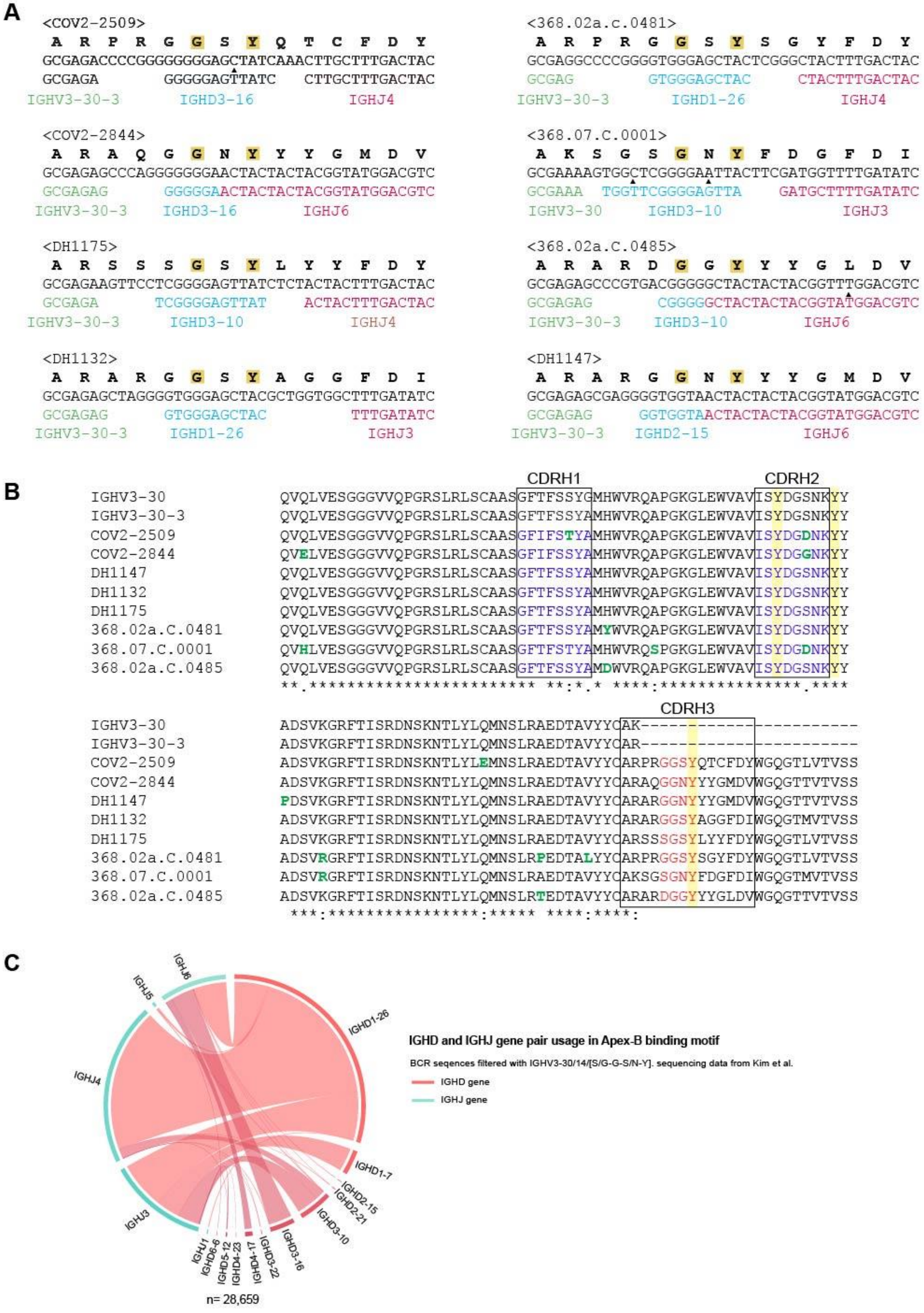
A
<COV2-2509>
A R P R G G S Y Q T C F D Y
GCGAGACCCCGGGGGGGGAGCTATCAAACTTGCTTTGACTAC
GCGAGA GGGGGAGTTATC CTTGCTTTGACTAC
IGHV3-30-3 IGHD3-16 IGHJ4
<368.02a.c.0481>
A R P R G G S Y S G Y F D Y
GCGAGGCCCCGGGGTGGGAGCTACTCGGGCTACTTTGACTAC
GCGAG GTGGGAGCTAC CTACTTTGACTAC
IGHV3-30-3 IGHD1-26 IGHJ4
<COV2-2844>
A R A Q G G N Y Y Y G M D V
GCGAGAGCCCAGGGGGGGAACTACTACTACGGTATGGACGTC
GCGAGAG GGGGGAACTACTACTACGGTATGGACGTC
IGHV3-30-3 IGHD3-16 IGHJ6
<368.07.C.0001>
A K S G S G N Y F D G F D I
GCGAAAAGTGGCTCGGGGAATTACTTCGATGGTTTTGATATC
GCGAAA TGGTTCGGGGAGTTA GATGCTTTTGATATC
IGHV3-30 IGHD3-10 IGHJ3
<DH1175>
A R S S S G S Y L Y Y F D Y
GCGAGAAGTTCCTCGGGGAGTTATCTCTACTACTTTGACTAC
GCGAGA TCGGGGAGTTAT ACTACTTTGACTAC
IGHV3-30-3 IGHD3-10 IGHJ4
<368.02a.C.0485>
A R A R D G G Y Y Y G L D V
GCGAGAGCCCGTGACGGGGGCTACTACTACGGTTTGGACGTC
GCGAGAG CGGGGGCTACTACTACGGTATGGACGTC
IGHV3-30-3 IGHD3-10 IGHJ6
<DH1132>
A R A R G G S Y A G G F D I
GCGAGAGCTAGGGGTGGGAGCTACGCTGGTGGCTTTGATATC
GCGAGAG GTGGGAGCTAC TTTGATATC
IGHV3-30-3 IGHD1-26 IGHJ3
<DH1147>
A R A R G G N Y Y Y G M D V
GCGAGAGCGAGGGGTGGTAACTACTACTACGGTATGGACGTC
GCGAGAG GGTGGTAACTACTACTACGGTATGGACGTC
IGHV3-30-3 IGHD2-15 IGHJ6
B
CDRH1
CDRH2
IGHV3-30 QVQLVESGGGVVQPGRSLRLSCAASGFTFSSYGMHWVRQAPGKGLEWVAVISYDGSNKYY
IGHV3-30-3 QVQLVESGGGVVQPGRSLRLSCAASGFTFSSYAMHWVRQAPGKGLEWVAVISYDGSNKYY
COV2-2509 QVQLVESGGGVVQPGRSLRLSCAASGFIFSTYAMHWVRQAPGKGLEWVAVISYDGDNKYY
COV2-2844 QVELVESGGGVVQPGRSLRLSCAASGFIFSSYAMHWVRQAPGKGLEWVAVISYDGGNKYY
DH1147 QVQLVESGGGVVQPGRSLRLSCAASGFTFSSYAMHWVRQAPGKGLEWVAVISYDGSNKYY
DH1132 QVQLVESGGGVVQPGRSLRLSCAASGFTFSSYAMHWVRQAPGKGLEWVAVISYDGSNKYY
DH1175 QVQLVESGGGVVQPGRSLRLSCAASGFTFSSYAMHWVRQAPGKGLEWVAVISYDGSNKYY
368.02a.C.0481 QVQLVESGGGVVQPGRSLRLSCAASGFTFSSYAMYWVRQAPGKGLEWVAVISYDGSNKYY
368.07.C.0001 QVHLVESGGGVVQPGRSLRLSCAASGFTFSTYAMHWVRQSPGKGLEWVAVISYDGDNKYY
368.02a.C.0485 QVQLVESGGGVVQPGRSLRLSCAASGFTFSSYAMDWVRQAPGKGLEWVAVISYDGSNKYY
CDRH3
IGHV3-30 ADSVKGRFTISRDNSKNTLYLQMNSLRAEDTAVYYCAK
IGHV3-30-3 ADSVKGRFTISRDNSKNTLYLQMNSLRAEDTAVYYCAR
COV2-2509 ADSVKGRFTISRDNSKNTLYLEMNSLRAEDTAVYYCARPRGGSYQTCFDYWGQGTLVTVSS
COV2-2844 ADSVKGRFTISRDNSKNTLYLQMNSLRAEDTAVYYCARAQGGNYYYGMDVWGQGTTVTVSS
DH1147 PDSVKGRFTISRDNSKNTLYLQMNSLRAEDTAVYYCARARGGNYYYGMDVWGQGTTVTVSS
DH1132 ADSVKGRFTISRDNSKNTLYLQMNSLRAEDTAVYYCARARGGSYAGGFDIWGQGTMVTVSS
DH1175 ADSVKGRFTISRDNSKNTLYLQMNSLRAEDTAVYYCARSSSGSYLYYFDYWGQGTLVTVSS
368.02a.C.0481 ADSVRGRFTISRDNSKNTLYLQMNSLRPEDTALYYCARPRGGSYSGYFDYWGQGTLVTVSS
368.07.C.0001 ADSVRGRFTISRDNSKNTLYLQMNSLRAEDTAVYYCAKSGSGNYFDGFDIWGQGTMVTVSS
368.02a.C.0485 ADSVKGRFTISRDNSKNTLYLQMNSLRTEDTAVYYCARARDGGYYYGLDVWGQGTTVTVSS
C
IGHJ6
IGHJ5
IGHD1-26
IGHJ4
IGHD1-7
IGHD2-15
IGHD2-21
IGHD3-10
IGHJ3
IGHJ1
IGHD6-6
IGHD5-12
IGHD4-23
IGHD4-17
IGHD3-22
IGHD3-16
n= 28,659
IGHD and IGHJ gene pair usage in Apex-B binding motif
BCR seqences filtered with IGHV3-30/14/[S/G-G-S/N-Y]. sequencing data from Kim et al.
IGHD gene
IGHJ gene

**Figure S7. Sequence analysis of the Apex-B epitope-directed antibodies.** (A) Amino acid and nucleotide sequences spanning the V-D-J junction are shown for eight Apex-B epitope-directed antibodies. Putative germline V, D, and J segments were assigned using IgBlast and are indicated below each sequence (V genes in green, D genes in blue, J genes in red). Conserved GxY motif residues in Apex-B antibodies are highlighted in yellow. (B) Amino acid sequence alignment of heavy-chain variable regions from the antibodies shown in (A), together with the germline sequence of IGHV3-30 and IGHV3-30-3. Within CDRH1-2, germline-conserved residues are highlighted in blue, whereas somatically mutated residues are shown in green. The G/S-G-S/N-Y motif within CDRH3 is highlighted in red. Three tyrosine residues mentioned in Figure 5B are colored in yellow boxes. Conserved residues are denoted by asterisks. (C) Chord diagram showing IGHD and IGHJ gene combinations identified from BCR sequence analysis of clones with Apex-B binding characteristics defined by IGHV3-30/14/[G/S-G-S/N-Y]

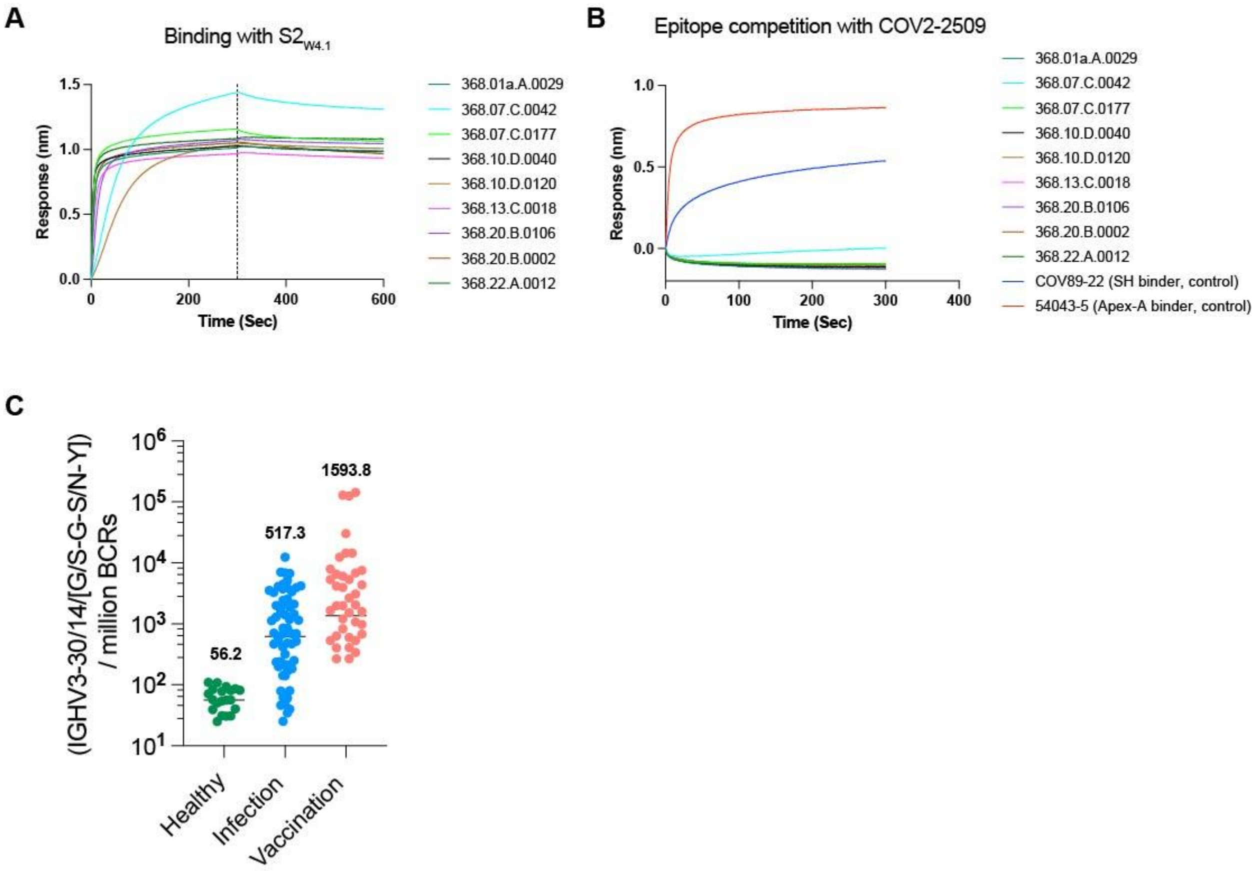


**Figure S8. Validation of Apex-B binding of clonally expanded IGHV3-30/14/Y-feature antibodies selected from individual donors in Figure 4J, and frequency analysis across donor groups with different exposure histories.** (A) Binding of most clonally expanded IGHV3-30/14/Y-feature antibodies to $S2_{W4.1}$, measured by biolayer interferometry (BLI). (B) Competitive BLI assay with COV2-2509, performed as described in Figure S5D. All donor-derived antibodies showed complete competition with COV2-2509, confirming targeting of the Apex-B epitope. COV89-22 (SH binder) and 54043-5 (Apex-A binder) were included as controls. (C) Frequency of IGHV3-30/14/Y-feature antibodies containing the G/S-G-S/N-Y motif in healthy, infected, and vaccinated cohorts, quantified by analysis of the OAS database and the Briney et al.[59]. repertoire datasets. Values are shown as the number of motif-containing BCRs per million sequences.

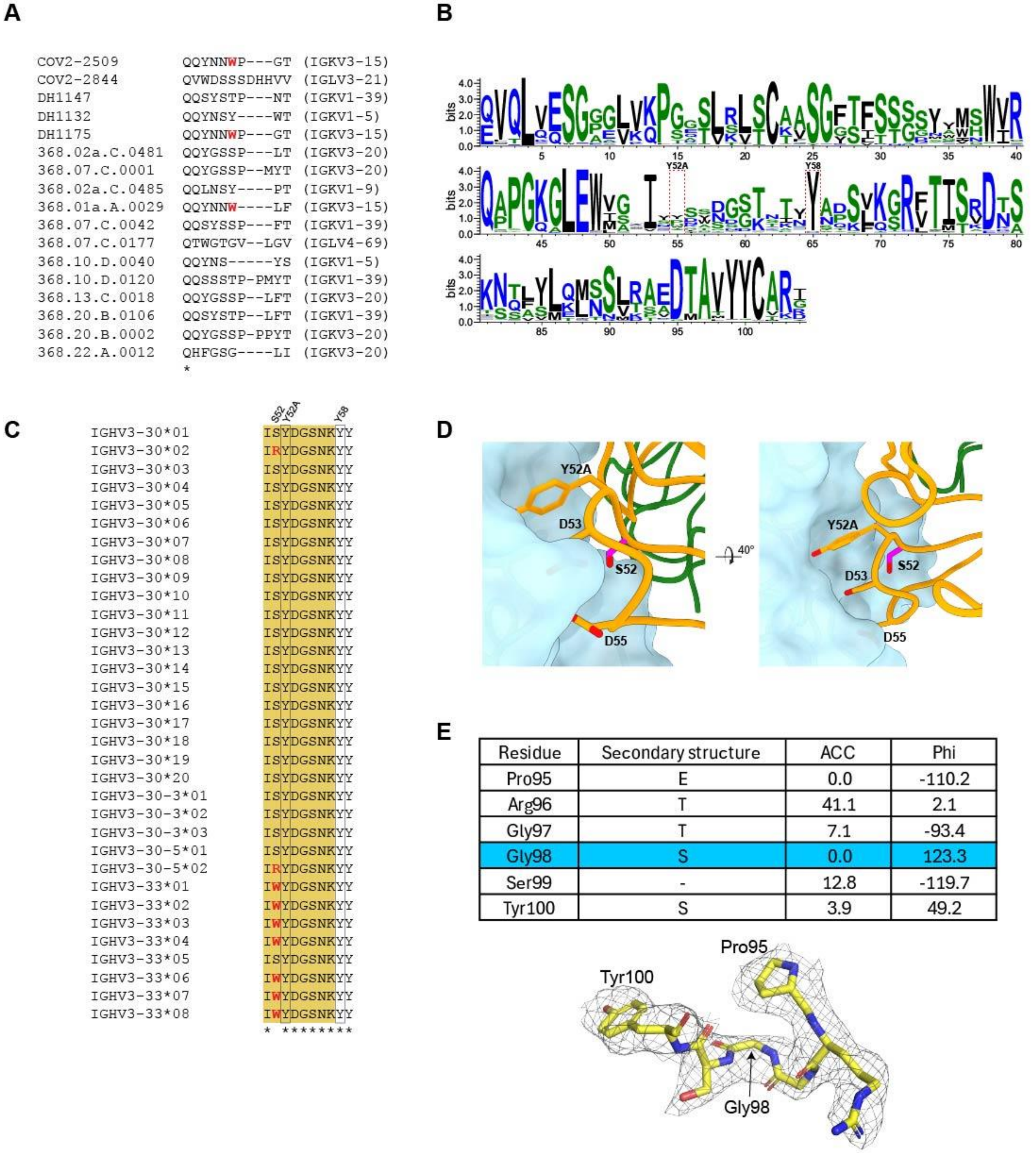


| Residue | Secondary structure | ACC | Phi |
|---|---|---|---|
| Pro95 | E | 0.0 | -110.2 |
| Arg96 | T | 41.1 | 2.1 |
| Gly97 | T | 7.1 | -93.4 |
| Gly98 | S | 0.0 | 123.3 |
| Ser99 | - | 12.8 | -119.7 |
| Tyr100 | S | 3.9 | 49.2 |

**Figure S9. Diverse light-chain usage, IGHV3-30-encoded tyrosine residue, and structural features in Apex B binding antibodies.** (A) Sequence alignment of light chains from Apex-B binding antibodies showing diversity in IGKV/IGLV gene usage and CDRL3 length (7-12 residues). The presence of Trp94 in CDRL3, encoded by IGKV3-15 in COV2-2509, is not conserved among Apex-B antibodies. (B) Sequence logo plots derived from 467 unmutated human IGHV germline reference sequences from the IMGT reference directory. The two tyrosine signature residues

(Tyr52A and Tyr58) characteristic of Apex-B-binding antibodies are indicated with dotted boxes. (C) Alignment of IGHV germlines that preserve both Tyr52A and Tyr58 residues. The two tyrosine residues are highly conserved within the IGHV3-30 subfamily and are highlighted with black boxes. The residues mutated at position Ser52 are shown in red bold, and the CDRH2 region is shaded in yellow. (D) Structural models based on the $S2_{W4.1}$-COV2-2509 complex structure show that substitutions at Ser52 (Trp or Arg) would cause steric clashes with surrounding residues, likely hindering optimal engagement with the Apex-B epitope. Ser52 is shown in pink, and the Apex-B epitope is displayed as a light-blue surface model, with the heavy and light chains of Fab colored yellow and green, respectively. (E) Backbone geometry and solvent accessibility in the G/S-G-S/N-Y motif region (CDRH3) of the COV2-2509 antibody. Secondary structure assignment, solvent accessibility, and backbone phi angles were derived using the DSSP tool[93]. Gly98, highlighted in cyan, displays an unusual positive phi angle and zero solvent accessibility, consistent with a structurally constrained and highly buried position. ACC denotes solvent accessible surface area as calculated by DSSP. Secondary structure assignments follow DSSP notation: E, beta strand; T, turn; and S, bend. Below, the EM density map corresponding to residues 95-100 of the CDRH3 is shown with the atomic model represented as sticks.

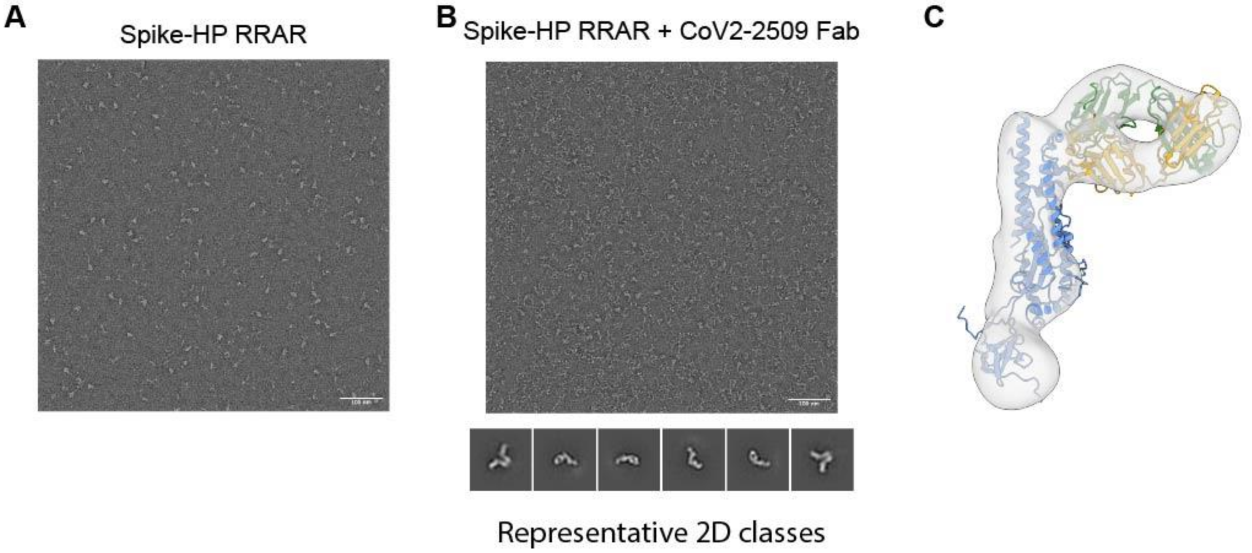


D

```
                737          750             765      845     853
                |            |               |        |       |
SARS-CoV-2      DCTMYICGDSTECSNLLLQYGSFCTQLNR         AARDLICAQ
SARS-CoV        DCNMYICGDSTECANLLLQYGSFCTQLNR         NARDLICAQ
GZ02            DCNMYICGDSTECANLLLQYGSFCTQLNR         NARDLICAQ
Rs4231          DCNMYICGDSTECANLLLQYGSFCTQLNR         NARDLICAQ
WIV1            DCNMYICGDSTECANLLLQYGSFCTQLNR         NARDLICAQ
YN2018C         DCTMYICGDSQECSNLLLQYGSFCTQLNR         NARDLICAQ
Rp3             DCTMYICGDSLECSNLLLQYGSFCTQLNR         SARDLICAQ
HKU3-1          DCTMYICGDSLECSNLLLQYGSFCTQLNR         SARDLICAQ
BM48-31         DCNMYICGDSTECSNLLLQYGSFCTQLNR         NARDLICAQ
RaTG13          DCTMYICGDSTECSNLLLQYGSFCTQLNR         AARDLICAQ
ZXC21           DCTMYICGDSIECSNLLLQYGSFCTQLNR         SARDLICAQ
ZC45            DCTMYICGDSIECSNLLLQYGSFCTQLNR         SARDLICAQ
Pangolin        DCTMYICGDSIECSNLLLQYGSFCTQLNR         AARDLICAQ
                **.******* **:***************          ********
```

**Figure S10. Structural disruption of spike trimer by COV2-2509 and Apex-B epitope conservation across sarbecoviruses.** (A) Ns-EM micrograph of Spike-HP RRAR alone. (B) ns-EM micrograph of Spike-HP RRAR incubated with COV2-2509 Fab, with representative 2D classes shown below. (C) 3D reconstruction of the ns-EM result shown in panel (B), fitted with the S2 monomer model in complex with COV2-2509 Fab from the $S2_{W4.1}$-COV2-2509 structure. (D) Sequence alignment of the Apex-B epitope region across sarbecoviruses. Conserved residues are denoted by asterisks.

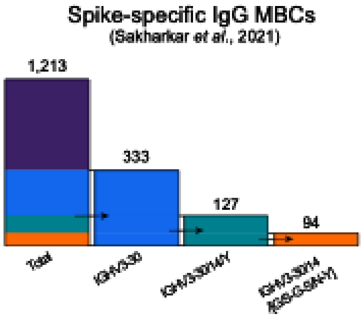


**Figure S11. Identification of Apex-B binding motif antibodies among spike-specific IgG MBCs.** Sequences from a total of 1,213 SARS-CoV-2 spike-specific IgG memory B cells were analyzed to identify antibodies harboring the Apex-B binding motif, using the monoclonal antibody sequence dataset reported by Sakharkar et al. (2021)[71].

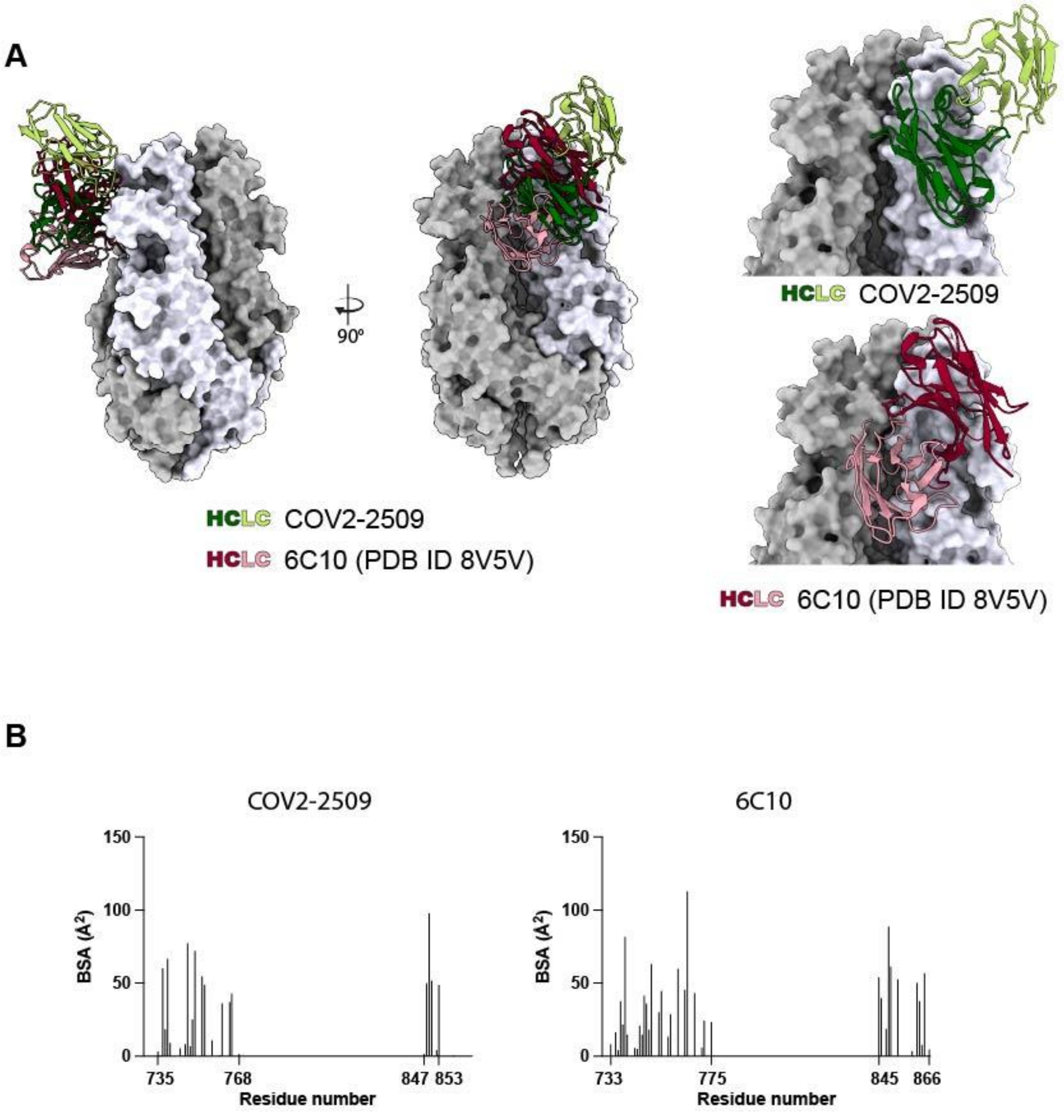


**Figure S12. Structural comparison of COV2-2509 and 6C10 binding to the S2 Apex-B region.** (A) Structural overlay of COV2-2509 and 6C10 (PDB ID 8V5V) Fabs bound to the S2 Apex-B epitope. The S2 trimer is shown as a surface representation, with the antibody-bound protomer highlighted in white and the remaining protomers in gray. Fabs are shown as cartoon models. Two orientations are shown, rotated 90°. The individual complex structures of COV2-2509 and 6C10 with S2 are shown in separate panels on the right, with all S2 structures derived from the COV2-2509-S2$_{W4.1}$ complex for consistency. (B) Buried surface area (BSA) analysis of antibody-antigen interfaces. COV2-2509 shows major contacts with residues 735-768 and 847-853, whereas 6C10 engages residues 733-775 and 845-866.

### Representative CryoEMPEM data processing workflow :SARS-CoV-2 S2 in complex with a pFab (Donor 3)

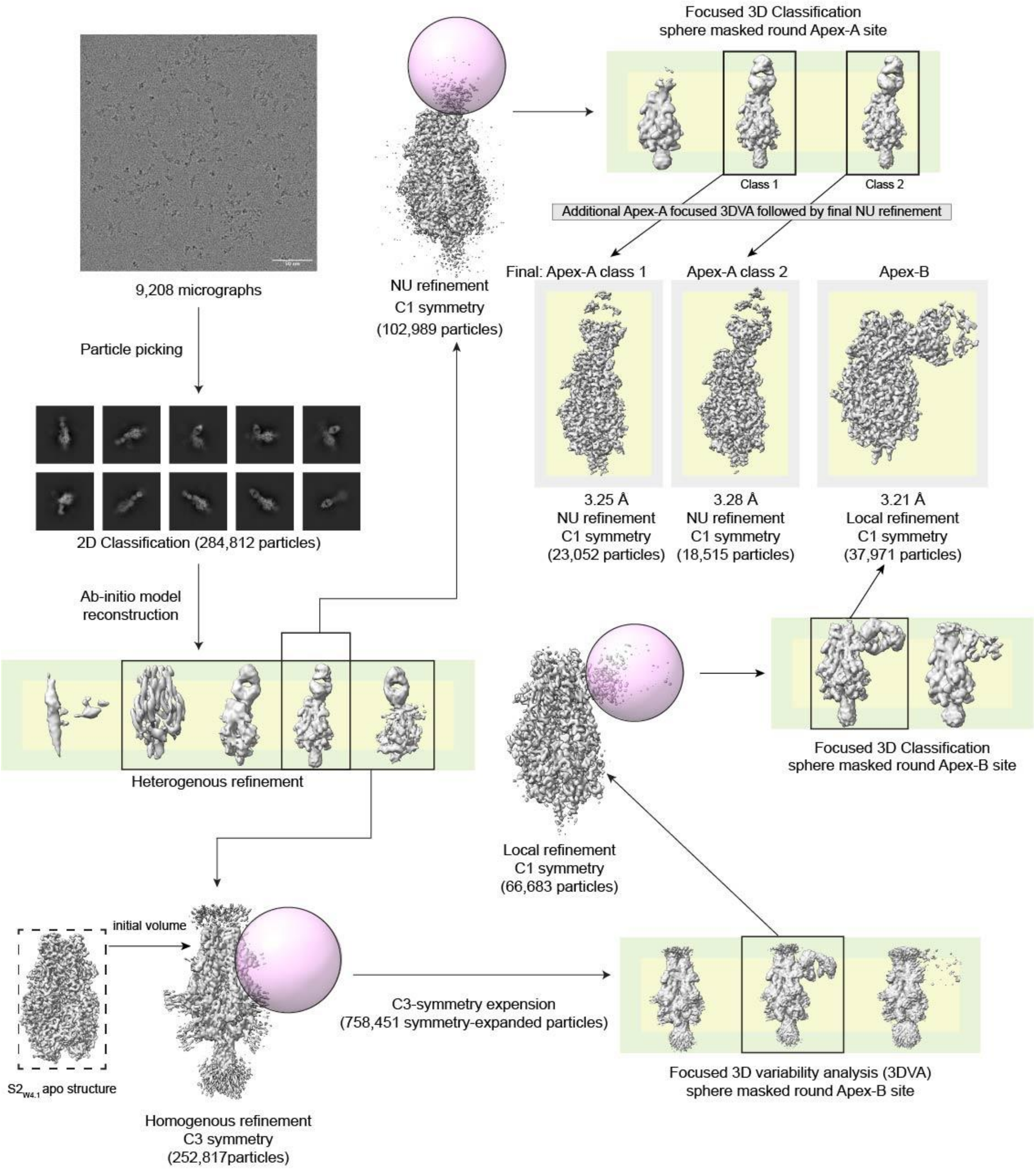


**Figure S13. Schematic representation of cryo-EMPEM data processing workflow.**

**Table S1. Stabilized SARS-CoV-2 S2 constructs**

| Protein name | Including residues | Introduced mutations | References |
|---|---|---|---|
| S2-E69 | 686-1208+foldon | Y707C-T883C, F970C-G999C, A892P, A899P, A942P, V987P, N907E, T961F, D994Q, Q1011M, I1018Y | Lee, J. et al., 2024[33] |
| S2-HP | 686-1208+foldon | F817P, A892P, A899P, A942P, K986P, V987P | Hsieh, C. L. et al., 2020[50] |
| S2-HP-2W | 698-1161 | S704C-K790C, F817P, A892P, A899P, A942P, K986P, V987P, Q957E, V991W, T998W | Nuqui, X. et al., 2024[34] |
| $S2_{W4}$ | 698-1161 | S704C-K790C, G880C-F888C, A892P, A899P, A942P, K986P, V987P, T961F | This study |
| $S2_{W4.1}$ | 698-1161 | S704C-K790C, G880C-F888C, A892P, A899P, A942P, K986P, V987P, T961F, V991W, T998W | This study |

*Mutated residues are colored by disulfide bonds (green), stabilizing prolines (blue), and trimer interface stabilizations (red).

**Table S2. Cryo-EM data collection, refinement, and validation statistics**

| | STS1 (Donor1) | STS2 (Donor2) | STS3 (Donor3) | STS4 (donor8-T2) | S2$_{W4.1}$ apo | S2$_{W4.1}$ +COV2-2509 Fab | Donor3 Apex-A class1 | Donor3 Apex-A class2 | Donor8-T2 Apex-A |
|---|---|---|---|---|---|---|---|---|---|
| | **EMD-73884** | **EMD-75295** | **EMD-74737** | **EMD-74738** | **EMD-74740** | **EMD-74739** | **EMD-74741** | **EMD-74742** | **EMD-74743** |
| | **PDB ID 9Z80** | **PDB ID 10MU** | **PDB ID 9ZT5** | **PDB ID 9ZT6** | **PDB ID 9ZT8** | **PDB ID 9ZT7** | **-** | **-** | **-** |
| **Data Collection and Processing** | | | | | | | | | |
| Electron microscope | TFS Glacios 2 | TFS Glacios 2 | TFS Glacios 2 | TFS Glacios 2 | TFS Glacios 2 | TFS Glacios 2 | TFS Glacios 2 | TFS Glacios 2 | TFS Glacios 2 |
| Electron detector | Falcon 4i | Falcon 4i | Falcon 4i | Falcon 4i | Falcon 4i | Falcon 4i | Falcon 4i | Falcon 4i | Falcon 4i |
| Magnification | 190,000x | 190,000x | 190,000x | 190,000x | 190,000x | 190,000x | 190,000x | 190,000x | 190,000x |
| Voltage (kV) | 200 | 200 | 200 | 200 | 200 | 200 | 200 | 200 | 200 |
| Electron exposure (e-/Å$^2$) | 45 | 45 | 45 | 45 | 45 | 45 | 45 | 45 | 45 |
| Defocus range (μm) | -0.8 to -1.8 | -0.8 to -1.8 | -0.8 to -1.8 | -0.8 to -1.8 | -0.8 to -1.8 | -0.8 to -1.8 | -0.8 to -1.8 | -0.8 to -1.8 | -0.8 to -1.8 |
| Pixel Size (Å) | 0.718 | 0.718 | 0.718 | 0.718 | 0.718 | 0.718 | 0.718 | 0.718 | 0.718 |
| Symmetry imposed | C1 | C1 | C1 | C1 | C3 | C1 | C1 | C1 | C1 |
| Number of movie micrographs | 8,375 | 13,189 | 15,610 | 7,672 | 5,579 | 2,763 | 15,610 | 15,610 | 7,672 |
| Number of particle images in map | 35,264 | 88,736 | 38,727 | 189,213 | 124,168 | 139,283 | 23,052 | 18,515 | 27,797 |
| Map resolution (Å) | 3.17 | 3.16 | 3.21 | 2.83 | 2.75 | 3.11 | 3.25 | 3.28 | 4.24 |
| FSC threshold | 0.143 | 0.143 | 0.143 | 0.143 | 0.143 | 0.143 | 0.143 | 0.143 | 0.143 |
| Map sharpening *B* factor (Å$^2$) | -50 | -60 | -49.6 | -55 | -52 | -72.7 | -49.7 | -46.9 | -95.9 |
| **Model building and validation** | | | | | | | | | |
| *Model composition* | | | | | | | | | |
| Protein Chains | 5 | 5 | 5 | 5 | 3 | 5 | - | - | - |

| | | | | | | | | | |
|---|---|---|---|---|---|---|---|---|---|
| Protein Residues | 1,446 | 1,454 | 1,454 | 1,446 | 1,278 | 1,617 | - | - | - |
| Glycans | 19 | | 21 | 24 | 24 | 24 | - | - | - |
| *RMSD bonds* | | | | | | | | | |
| Bond lengths (Å) | 0.003 | 0.003 | 0.004 | 0.004 | 0.003 | 0.005 | - | - | - |
| Bond angles (°) | 0.65 | 0.63 | 0.69 | 0.70 | 0.56 | 1.01 | - | - | - |
| *Ramachandran plot* | | | | | | | | | |
| Favored (%) | 96.86 | 98.30 | 96.60 | 97.79 | 97.84 | 97.57 | - | - | - |
| Outliers (%) | 0 | 0 | 0 | 0 | 0 | 0 | - | - | - |
| *Validation* | | | | | | | | | |
| MolProbity score | 1.41 | 1.11 | 1.51 | 1.22 | 1.17 | 1.20 | - | - | - |
| Clashscore | 4.40 | 3.23 | 5.41 | 3.83 | 3.46 | 3.16 | - | - | - |
| Rotamer outliers (%) | 0.07 | 0.19 | 0.28 | 0 | 0 | 0.14 | - | - | - |
| EMRinger score | 3.24 | 3.44 | 2.43 | 3.86 | 4.48 | 3.78 | - | - | - |
| Map-model correlation coefficient, CC (mask) | 0.83 | 0.84 | 0.80 | 0.84 | 0.90 | 0.79 | - | - | - |
| CaBLAM outliers (%) | 2.44 | 1.54 | 2.19 | 1.40 | 1.97 | 1.8 | - | - | - |

**Table S3. Demographics of study donors**

| | Sample ID | Sex | Age bin | SARS-CoV-2 immune history | Year of last immune event |
|---|---|---|---|---|---|
| Donor1 | C0751 | male | 18-29 | M-M-M | 2021 |
| Donor2 | A2173 | male | 30-39 | P-P-P | 2021 |
| Donor3 | C2173 | female | 18-29 | M-M-M | 2022 |
| Donor4 | C8241 | female | 18-29 | I-P-P-P-I-$M^b$ | 2022 |
| Donor5 | A6865 | female | 18-29 | M-M-P-I | 2022 |
| Donor6 | A8903 | female | 50-59 | I | 2022 |
| Donor7 | C0515 | female | 30-39 | I-I | N/A |
| Donor8-T1 | A0230 | female | 40-49 | P-P-P | 2021 |
| Donor8-T2 | A3500 | female | 40-49 | P-P-P | 2022 |
| Donor8-T3 | C0279 | female | 40-49 | P-P-P-$M^b$/$P^b$-I | 2023 |
| Donor9-T1 | A4922 | female | 50-59 | P-P-P-I | 2021 |
| Donor9-T2 | A5538 | female | 50-59 | P-P-P-I | 2022 |

M: Moderna (mRNA-1273) vaccine; P: Pfizer (BNT162b2) vaccine; I: infection. [b]Bivalent vaccine, $M^b$/$P^b$: bivalent vaccine, unclear whether Moderna or Pfizer. N/A: not available.

**Table S4. Antibody titer against SARS-CoV-2 spike in each donor**

| Donor | Spike-binding (endpoint titer) | Spike-binding (AUC) |
|---|---|---|
| Donor 1 | 12,800 | 3,496 |
| Donor 2 | 12,800 | 4,063 |
| Donor 3 | 6,400 | 3,151 |
| Donor 4 | 6,400 | 3,041 |
| Donor 5 | 102,400 | 24,421 |
| Donor 6 | 3,200 | 1,095 |
| Donor 8-T1 | 1,600 | 379.7 |
| Donor 8-T2 | 12,800 | 6,062 |
| Donor 8-T3 | 12,800 | 4,119 |
| Donor 9-T1 | 25,600 | 5,091 |
| Donor 9-T2 | 12,800 | 5,505 |

**Table S5. References for antibodies with 8th tyrosine and 14-residue of CDRH3 identified in the CoV-AbDab database**

| Reference | 14 length and 8th Y CDRH3 antibodies (count) |
|---|---|
| Wheatley, A.K. *et al*., 2021[94] | 5 |
| Shiakolas, A.R. *et al*., 2021[95] | 1 |
| Shiakolas, A.R. *et al*., 2022[96] | 1 |
| Wec, A.Z. *et al*., 2020[97] | 4 |
| Graham, C. *et al*., 2021[98] | 9 |
| Gaebler, C. *et al*., 2021[99] | 2 |
| Li, D. *et al*., 2021[55] | 39 |
| Robbiani, D.F. *et al*., 2020[46] | 1 |
| Pinto, D. *et al*., 2021[11] | 3 |
| Seydoux, E. *et al*., 2020[100] | 6 |
| Song, G. *et al*., 2021[74] | 1 |
| Turner, J. S. *et al*., 2021[101] | 1 |
| Scheid, J. F. *et al*., 2021[102] | 1 |
| Vanshylla, K. *et al*., 2022[103] | 18 |
| Kramer, K. J. *et al*., 2022[104] | 2 |
| Huang, K. A. *et al*., 2023[105] | 1 |
| Sakharkar, M. *et al*., 2021[71] | 126 |
| Tong, P. *et al*., 2021[106] | 2 |
| Brouwer, P. J. M. *et al*., 2020[107] | 2 |
| Zost, S. J. *et al*., 2020[54] | 20 |
| Rogers, T. F. *et al*., 2020[108] | 1 |
| He, W. T. *et al*., 2022[109] | 1 |
| Dejnirattisai, W.*et al*., 2021[110] | 1 |
| Kim, W *et al*., 2022[53] | 211 |
| Chi, X., *et al*., 2020[111] | 1 |
| Zhou, Y., *et al*., 2021[112] | 3 |
| Cao, Y., *et al*., 2022[113] | 1 |
